# Atomic resolution map of the solvent interactions driving SOD1 unfolding in CAPRIN1 condensates

**DOI:** 10.1101/2024.04.29.591724

**Authors:** Rashik Ahmed, Mingyang Liang, Rhea P. Hudson, Atul K. Rangadurai, Shuya Kate Huang, Julie D. Forman-Kay, Lewis E. Kay

**Affiliations:** Department of Molecular Genetics, University of Toronto, Toronto, ON, Canada, M5S 1A8; Department of Chemistry, University of Toronto, Toronto, ON, Canada, M5S 3H6; Department of Biochemistry, University of Toronto, Toronto, ON, Canada, M5S 1A8; Program in Molecular Medicine, Hospital for Sick Children Research Institute, Toronto, ON, Canada, M5G 0A4

**Keywords:** methyl-TROSY NMR, phase separation, folding-unfolding equilibria, protein stability, client/scaffold

## Abstract

Biomolecules can be sequestered into membrane-less compartments, referred to as biomolecular condensates. Experimental and computational methods have helped define the physical-chemical properties of condensates. Less is known about how the high macromolecule concentrations in condensed phases contribute “solvent” interactions that can remodel the free-energy landscape of other condensate-resident proteins, altering thermally accessible conformations and, in turn, modulating function. Here, we use solution Nuclear Magnetic Resonance (NMR) spectroscopy to obtain atomic resolution insights into the interactions between the immature form of superoxide dismutase 1 (SOD1), which can mislocalize and aggregate in stress granules, and the RNA-binding protein CAPRIN1, a component of stress granules. NMR studies of CAPRIN1:SOD1, focused on both unfolded and folded SOD1 states in mixed phase and de-mixed CAPRIN1-based condensates, establish that CAPRIN1 shifts the folding equilibrium of SOD1 towards the unfolded state through preferential interactions with the unfolded ensemble, with little change to the structure of the folded conformation. Key contacts between CAPRIN1 and the H80-H120 region of unfolded SOD1 are identified, as well as SOD1 interaction sites near both the arginine-rich and aromatic-rich regions of CAPRIN1. Unfolding of immature SOD1 in the CAPRIN1 condensed phase is shown to be coupled to aggregation, while a more stable zinc-bound, dimeric form of SOD1 is less susceptible to unfolding when solvated by CAPRIN1. Our work underscores the impact of the condensate solvent environment on the conformational states of resident proteins and supports the hypothesis that ALS mutations that decrease metal binding or dimerization function as drivers of aggregation in condensates.

**Significance Statement:** Biomolecular condensates concentrate proteins and nucleic acids to regulate and perform key biological functions. Although the material properties of these condensates are well-studied, much less is understood about how the structure and dynamics of proteins within them are affected by the high concentration of biomolecules. In this study we have used NMR spectroscopy to study how the folding equilibrium and structural dynamics of the ALS protein SOD1 are modulated inside a condensate formed by CAPRIN1. Our study reveals that the CAPRIN1 condensed phase biases an immature form of SOD1 towards unfolded states that are susceptible to aggregation and provides insights into why this is the case, while a more mature form of the protein is much less affected.

## Introduction

The compartmentalization of biomolecules into distinct membrane-bound organelles is critical for the specificity and fidelity of biochemical processes. Beyond this well-known mechanism of organization, biomolecules are also sorted into membrane-less compartments, referred to as biomolecular condensates, understood to form through the process of phase separation (1, 2). Condensates, formed through multivalent interactions between “scaffold” molecules, create unique solvent environments that enrich a subset of the cellular milieu (“clients”) while excluding others (1– 3). Through such compositional control, condensates can enhance (inhibit) enzyme activity, modulate specificity of reactions, and control substrate binding (2, 4). Such concentration-dependent (*i.e*., mass action) mechanisms of reaction rate changes in condensates have been described in diverse systems (5–8). Accordingly, modifying condensate composition through post-translational modifications (PTMs) and changes in environmental conditions offers a versatile and dynamic approach to regulating biochemical reactions (9).

Concentration-independent mechanisms of enzymatic activity changes have also been described. The distinct material properties of condensates, which are highly viscous compared to the surrounding dilute phase (10–12), can modulate reaction rates through changes in binding kinetics, k_on_ and k_off_, and by limiting the diffusion of enzymatic reaction components away from sites of activity (increasing dwell time) (13, 14). Moreover, the condensate solvent environment is distinct from the surrounding dilute phase. The concentrations of proteins in condensed phases of phase-separated systems can exceed tens of millimolar (15–17), with a concomitant decrease in the water content to as low as 65% of the dilute phase (10, 18). Under these conditions, the scaffolding proteins significantly contribute to the solvation of the resident molecules. Such proteinaceous solvents exhibit properties akin to organic solutions, with estimates of dielectric constants and polarity indicating similarities to DMSO (11) and methanol (19), respectively. Thus, condensate solvent environments are hypothesized to modulate the conformational equilibria of client proteins, as has already been demonstrated for nucleic acids (20–23). However, insights into the conformational changes of proteins inside condensed phases have thus far been inferred based on enhancements of enzymatic activities that are not fully accounted for by mass action (6, 24, 25). In some cases they have been directly observed, albeit at low resolution (26), precluding meaningful insights into structure-function relationships inside condensates.

Here, we address this gap by directly probing the conformational changes of a client protein inside a model scaffold condensate at atomic resolution through solution NMR (Fig. 1A). Our system consists of the client metalloenzyme superoxide dismutase 1 (SOD1), which catalyzes the disproportionation of superoxide radicals into oxygen and hydrogen peroxide (27). The nascent metal-free, disulfide-reduced form of the enzyme exists as a monomer that exchanges between an unfolded minor state and a folded major conformer (Fig. 1B, *left*). This immature form of the enzyme has a high propensity to misfold and aggregate *in vitro* (28, 29). In contrast, the fully mature state forms a stable homodimer, with each SOD1 subunit bound to a copper and a zinc ion, and an intra-subunit disulfide bridge formed between C57 and C146 (Fig. 1B, *right*) (29). Mutations in the *sod1* gene account for ~20% of familial amyotrophic lateral sclerosis (ALS) cases and 2% of all ALS occurrences (30), and enhance the accumulation of insoluble neuronal inclusions containing SOD1 (31). Notably, such mutations also increase the mislocalization of SOD1 into stress granules (32) containing the RNA-binding protein CAPRIN1, among others.

**Fig. 1.**
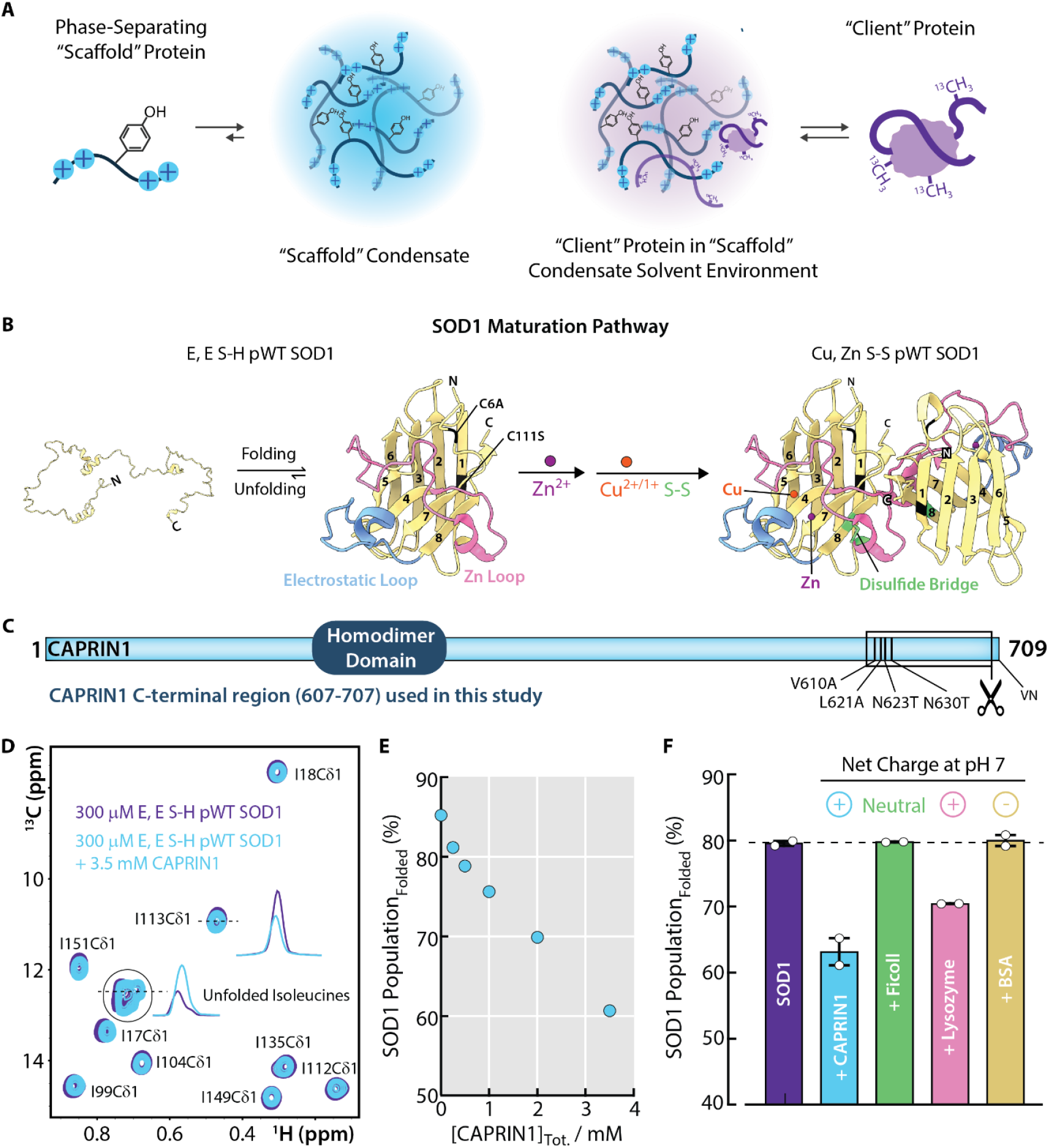
Stress granule “scaffold” protein CAPRIN1 solvates the SOD1 “client” and biases it towards the unfolded ensemble. (A) Schematic depicting the solvation of a “client” protein in a “scaffold” protein condensate. (B) Maturation pathway of SOD1. In the absence of metals and disulfide bond formation involving C146 and C57, E, E S-H SOD1 is in equilibrium between an unfolded ensemble and a folded conformation consisting of an 8-stranded β-barrel with two long loops. Upon Cu, Zn metalation and intra-subunit disulfide bond formation, Cu, Zn S-S SOD1 adopts a homodimeric state. The pseudo-WT (pWT) construct used in this study, with cysteine mutants C6A and C111S color-coded in black on the SOD1 structure, is shown (36) (PBD ID: 1HL5). The electrostatic loop and the zinc (dimer) loop are shown in blue and pink, respectively, the intra-subunit disulfide bridge is shown in green, and the zinc and copper metals are depicted as purple and orange spheres, respectively. (C) Schematic of the construct and associated mutations/truncation of CAPRIN1 used in this study. (D) Selected regions of [^1^H, ^13^C] HMQC spectra of the methyl region of 300 µM E, E S-H pWT SOD1 in the absence (*purple*) and presence (*blue*) of 3.5 mM CAPRIN1, 25 °C, 800 MHz. 1D slices of representative peaks are shown, highlighting the shift in population of folded and unfolded conformers upon addition of CAPRIN1. (E) Population of SOD1 folded conformation as a function of CAPRIN1 concentration, 25 °C. Population fractions were calculated as 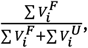 where 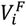 is the volume of folded (F) peak *i*, and 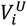 the volume of unfolded (U) peak *i*. The volume of the unfolded isoleucine cluster was determined through a sum-over-box approach. (F) Population of SOD1 folded conformation in the presence of various crowders, based on the integrals of ^19^F resonances of fluorotryptophan-labeled SOD1 derived from folded and unfolded conformers shown in *SI Appendix*, Fig. S1A,D-G. Error bars report standard deviations of replicates (circles). Crowders are annotated based on their net charge at pH 7, either neutral, positive (+), or negative (-).

In this study we have used the C-terminal low complexity region (607-707) of CAPRIN1 (this 101 residue fragment is referred to in what follows as CAPRIN1), which is sufficient to undergo phase separation (15), as a scaffold to solubilize the immature form of SOD1 (Fig. 1C) so that its properties can be studied in a highly proteinaceous condensed phase solvent environment using solution NMR spectroscopy. In particular, NMR experiments performed in a scaffold condensed phase and in mixed solutions with high scaffold concentrations reveal a shift in the conformational equilibria of the nascent protein towards the unfolded ensemble. We show that the conformational change arises from the preferential solvation of the unfolded ensemble by CAPRIN1, with interaction sites broadly distributed across the CAPRIN1 sequence, yet predominantly localized to a region spanning residues ~80-120 of unfolded SOD1. The unfolding is partially mimicked by proteinaceous solvents with positive charge, as for CAPRIN1, but not by the negatively charged bovine serum albumin (BSA) protein. Despite comparable SOD1 concentrations in condensed and dilute phases of CAPRIN1, SOD1 aggregation is exclusively observed in the condensed phase, highlighting a concentration-independent mechanism by which condensates may accelerate the progression of neurodegenerative disorders, such as ALS. Notably, the more stable zinc-bound, dimeric form of SOD1 (melting temperature 10 °C higher relative to the immature state (33–35)) has increased resilience to unfolding, underscoring the role of ALS mutations that either weaken metal binding or disrupt dimerization as potential drivers of pathogenesis in stress granules. Collectively, these findings highlight the role of scaffold proteins as solvents of client molecules in condensates, and suggest a mechanism by which cells can dynamically alter condensate composition, and accordingly their solvent nature, to modulate the conformational states of resident proteins and thereby achieve specific biological functions.

## Results

### CAPRIN1 solvation biases immature SOD1 towards an unfolded ensemble

To evaluate the impact of a scaffold condensate on the conformational equilibria of the client nascent SOD1, we first mimicked the condensed phase solvent environment using high concentration mixed solutions of the CAPRIN1 scaffold (*i.e*., not phase-separated). In the present study we have used the N623T/N630T mutant of the C-terminal region of CAPRIN1 as this double mutation eliminates formation of iso-Asp linkages which have been previously observed to form over time with the wild-type sequence (12). Further, in order to avoid the possibility of spectral overlap between methyl resonances of CAPRIN1 and those from the much less concentrated nascent SOD1 client, we have made the following additional mutations in CAPRIN1: V610A/L621A, with elimination of the C-terminal residues V708 and N709. None of these mutations affected the propensity of CAPRIN1 to phase separate. In a similar manner, the non-conserved cysteines, C6 and C111, of SOD1 have been replaced with Ala and Ser (36), respectively, to prevent intermolecular disulfide bond formation during the course of our experiments. This SOD1 variant is referred to as pseudo wild-type (pWT) SOD1 in what follows (37).

Heteronuclear multiple quantum correlation (HMQC) spectra of the disease-relevant, immature pWT SOD1 form (denoted as E, E S-H pWT SOD1 to indicate that both Cu and Zn metal binding sites of the monomer are empty “E” and that C57 and C146 are reduced) were recorded on samples with ^2^H, ^15^N,^13^C-ILV methyl labeling, both in the absence and presence of 3.5 mM unlabeled CAPRIN1 (Fig. 1D). In this labeling scheme only methyl groups of Ile (δ1), Leu(δ1,δ2), and Val(γ1,γ2) are NMR observable (^13^CH_3_), with only one of the two prochiral methyl groups of Leu and Val ^13^CH_3_-labeled (38). Under these conditions, E, E S-H pWT SOD1 exists as a monomer in slow-exchange on the NMR chemical shift timescale between a folded major conformation and an unfolded minor state, giving rise to two separate sets of NMR resonances (39–41). Correlations in [^1^H,^13^C] HMQC spectra arising from the SOD1 isoleucine methyl groups provide a straightforward readout of this folding-unfolding equilibrium, with nine well-dispersed resonances from the folded conformer and a corresponding set of overlapping signals from the unfolded ensemble (Fig. 1D). Upon addition of 3.5 mM CAPRIN1, a dramatic reduction in peak volumes from the folded conformer is observed with a concomitant enhancement in the volumes of the unfolded signals, indicating a shift of the equilibrium towards the unfolded ensemble (Fig. 1D, *blue vs. purple*). A titration of CAPRIN1, from 0 – 3.5 mM, into a solution of E, E S-H pWT SOD1 reveals a dose-dependent loss of the SOD1 folded population from 85% to 61% (Fig. 1E). Notably, the SOD1 unfolding profile is not saturable at CAPRIN1 concentrations up to 3.5 mM (Fig. 1E), indicative of weak “solvent” (CAPRIN1) interactions with SOD1.

Next, we evaluated whether the changes in the E, E S-H pWT SOD1 equilibrium observed upon addition of CAPRIN1 are reproduced when other proteins or the inert sugar polymer ficoll are added at high concentrations (43 mg/mL was used here). We recorded ^19^F NMR spectra of E, E S-H pWT SOD1 with the endogenous tryptophan in β-strand 3 substituted with 5-fluorotryptophan, both in the absence and presence of various crowders (*SI Appendix*, Fig. S1). Two resonances, corresponding to the folded and unfolded states, were observed and unambiguously assigned on the basis of high salt and high temperature conditions which stabilize the unfolded ensemble (*SI Appendix*, Fig. S1A-C). Addition of CAPRIN1 at high concentration confirms the unfolding phenomenon observed in HMQC spectra (Fig. 1F, *blue vs. purple*; *SI Appendix*, Fig. S1D *vs*. S1A). In contrast, addition of ficoll did not alter the SOD1 folding equilibrium (Fig. 1F, *green vs. purple*; *SI Appendix*, Fig. S1E *vs*. S1A). To determine whether other proteinaceous solvents can recapitulate the effect observed with CAPRIN1, we also recorded ^19^F NMR spectra in the presence of lysozyme, similarly positively charged as CAPRIN1 (+9 lysozyme *vs*. +13 CAPRIN1), and negatively-charged BSA (−13; *SI Appendix*, Fig. S1F-G). Whereas lysozyme partially unfolds SOD1, no significant change is observed in the presence of BSA (Fig. 1F, *pink and yellow vs. purple*; *SI Appendix* Fig. S1F,G *vs*. S1A). As E, E S-H pWT SOD1 is negatively charged (−6), our findings suggest that a contributor to its unfolding is electrostatic interactions with the proteinaceous solvent.

### Intermolecular CAPRIN1 : E, E S-H pWT SOD1 interactions

In order to gain insight into why addition of CAPRIN1 leads to a shift in the E, E S-H pWT SOD1 folding equilibrium towards the unfolded ensemble, we performed intermolecular PRE experiments which quantify interactions between proximal CAPRIN1 and SOD1 molecules on a per-residue basis. Cysteines were introduced, one at a time, into an otherwise cysteine-less CAPRIN1 construct to produce S615C, A658C or S678C variants and a 1,4,7,10-tetraazacyclododecane-1,4,7,10-tetraacetic acid (DOTA) cage loaded with either gadolinium (Gd^3+^, paramagnetic) or lutetium (Lu^3+^, diamagnetic) was conjugated on the incorporated cysteine through a maleimide linkage (Fig. 2A). The metal-bound cages did not compromise homotypic CAPRIN1 interactions, as confirmed by comparable phase-separation propensities with the non-caged construct (*SI Appendix*, Fig. S2), and the cage-conjugated CAPRIN1 molecules are therefore expected to exhibit largely unperturbed heterotypic interactions with SOD1 as well, as we will subsequently show. ^15^N-labeled E, E S-H pWT SOD1 was mixed with sub-stoichiometric amounts of either paramagnetic or diamagnetic ^14^N-CAPRIN1 (in a 3 SOD1:1 CAPRIN1 ratio) and intermolecular interactions were read-out via the intensities of peaks in fully relaxed [^1^H,^15^N]-TROSY HSQC spectra (42).

**Fig. 2.**
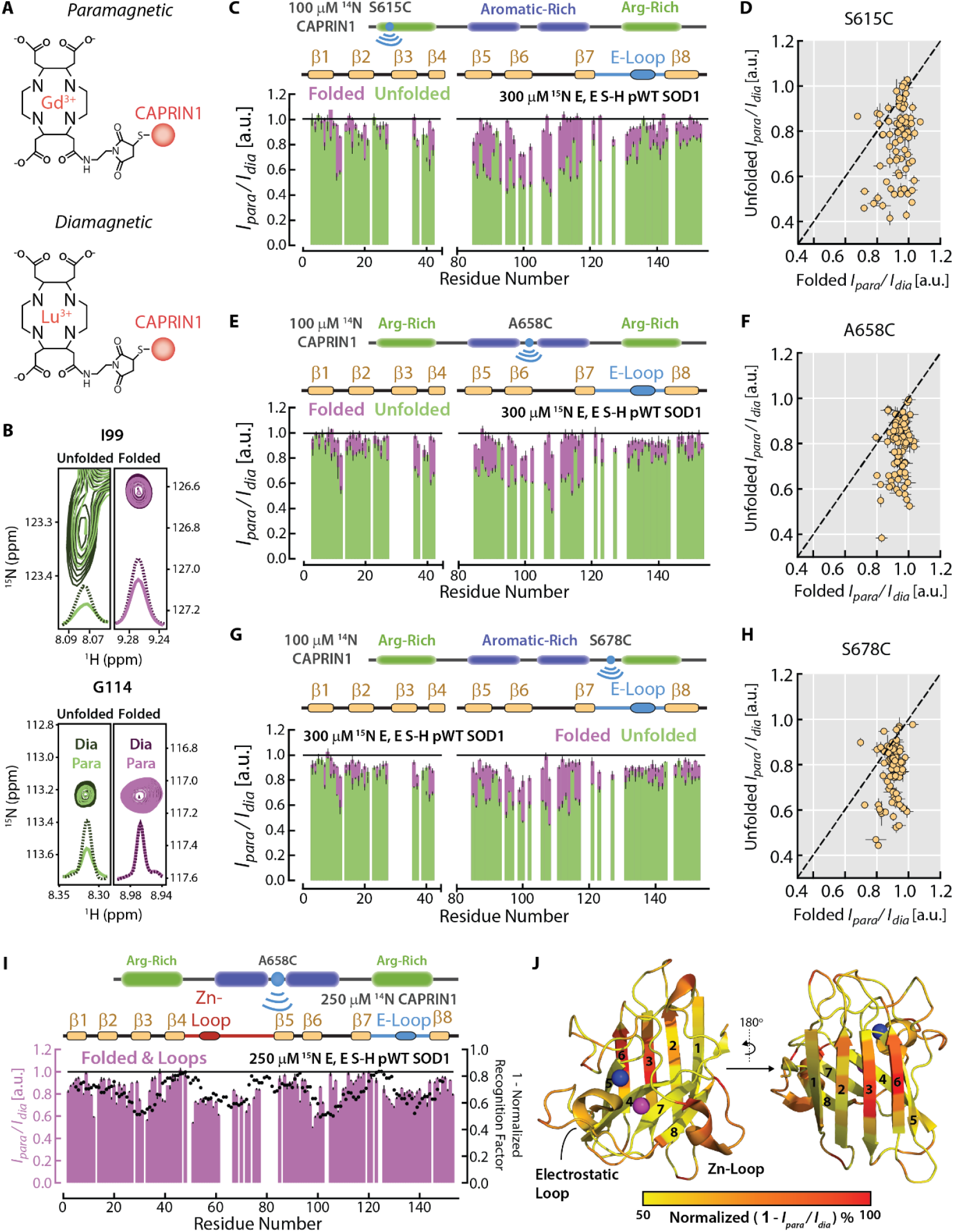
Intermolecular interactions of CAPRIN1 with folded and unfolded E, E S-H pWT SOD1. (A) Cartoon depicting the conjugation of a maleimide DOTA cage onto an exogenously introduced Caprin1 cysteine. Cages are filled with either paramagnetic (gadolinium) or diamagnetic (lutetium) metal. (B) Overlay of selected slices through peaks derived from unfolded (*green*) and folded (*purple*) E, E S-H pWT SOD1 for two representative residues, I99 and G114, in diamagnetic (*darker colors, dashed lines)* and paramagnetic (*lighter colors, solid lines*; spin-label at position 615) spectra. (C) Intensity profiles, *I*_*para*_/*I*_*dia*_, for 300 µM ^15^N E, E S-H pWT SOD1 in the presence of 100 µM ^14^N CAPRIN1 S615C-maleimide-DOTA coordinated with either gadolinium (*I*_*para*_) or lutetium (*I*_*dia*_). Error bars report standard deviations of replicates (*SI Appendix*, Materials and Methods). Intermolecular *I*_*para*_/*I*_*dia*_ ratios measured for folded (unfolded) E, E S-H pWT SOD1 are shown in purple (green). Only residues with unique folded and unfolded resonances are shown. A schematic diagram of CAPRIN1, with arginine-rich (*green rectangles*) and aromatic-rich (*purple rectangles*) regions highlighted along with the cysteine mutation for the maleimide-DOTA cage conjugation site (*blue circle*), and the secondary structure elements of folded SOD1, are displayed above the intermolecular PRE profile. Electrostatic loop is abbreviated as (E)-loop. (D) Linear correlation plot of *I*_*para*_/*I*_*dia*_ ratios for folded and unfolded E, E S-H pWT SOD1. Dashed line is y=x. (E)-(H) As (C)-(D), except with 100 µM ^14^N CAPRIN1 A658C-maleimide-DOTA (E)-(F) or 100 µM ^14^N CAPRIN1 S678C-maleimide-DOTA (G)-(H). (I) Intermolecular *I*_*para*_/*I*_*dia*_ profiles for 250 µM ^15^N E, E S-H pWT SOD1 in the presence of 250 µM ^14^N CAPRIN1 A658C-maleimide-DOTA (*purple bars*). Values derived from all non-overlapping folded SOD1 resonances and resonances from loop regions with identical folded and unfolded chemical shifts are shown. The recognition factor, an aggregate measure of the hydrophilicity and average interaction energy of an amino acid (44), is computed for the SOD1 amino acid sequence, normalized, averaged over a 9-residue window and plotted as 1 – Normalized Recognition Factor (*black circles*). (J) Heat map of CAPRIN1 interaction sites in folded SOD1 plotted onto the structure of a single protomer from the mature SOD1 structure (PDB ID: 1HL5) (36). The relative strengths of the intermolecular interactions, used for the heat map, were calculated as 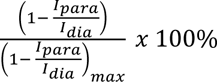, based on the *I*_*para*_/*I*_*dia*_ profile shown in (I). All NMR experiments were recorded at 25 °C, 800 MHz.

Fig. 2B highlights a number of regions from HSQC spectra showing differences in peak intensities (illustrated through cross sections) in spectra recorded with either Gd^3+^- or Lu^3+^-DOTA tagged CAPRIN1. As Gd^3+^ has seven unpaired f-electrons for an electron spin quantum number of 7/2, while Lu^3+^ is spin 0, the large magnetic field produced by the paramagnetic Gd^3+^ significantly attenuates proximal ^1^H probes on E, E S-H pWT SOD1 in a manner that depends on the metal-^1^H distance and the lifetime of the Gd^3+^-^1^H interaction. Thus, peaks in the Gd-DOTA tagged CAPRIN1 spectrum will be attenuated relative to the corresponding peaks in the Lu-DOTA dataset. This “so-called” paramagnetic relaxation enhancement (PRE) effect is most simply quantified in terms of relative peak intensities of SOD1 resonances in spectra recorded with paramagnetic *vs*. diamagnetic CAPRIN1 (*I*_*para*_ */ I*_*dia*_). As separate sets of peaks are observed for folded and unfolded E, E S-H pWT SOD1, the magnitude of CAPRIN1 interactions with each state can be probed separately (Fig. 2B Unfolded *I*_*para*_ */ I*_*dia*_, *green* and Folded *I*_*para*_ */ I*_*dia*_, *purple*), as illustrated for I99 and G114. While *I*_*para*_ */ I*_*dia*_ < 1 for the I99 amide proton in both unfolded and folded states, only in the unfolded state is there an appreciable reduction in *I*_*para*_ */ I*_*dia*_ for G114. Thus, the region of CAPRIN1 containing the DOTA-tag (S615C) is distal from the amide proton of G144 in the folded conformer of E, E S-H pWT SOD1 and/or has a relative smaller lifetime of interaction.

Experiments using CAPRIN1 spin-labeled in the N-terminal arginine-rich region (S615C) revealed, on average, a dramatic reduction in the *I*_*para*_ */ I*_*dia*_ ratios for unfolded E, E S-H pWT SOD1 compared to the folded conformer (Fig. 2C, *green vs. purple*, Fig. 2D), indicating preferential solvation of the unfolded ensemble by CAPRIN1. Such interaction bias explains the shift in the folding equilibrium towards the unfolded state, which is stabilized by CAPRIN1. The CAPRIN1 interaction site in unfolded E, E S-H pWT SOD1 is primarily localized to a region spanning residues ~ 80-120 (Fig. 2C, *green*). To establish whether there are additional SOD1 interaction sites in CAPRIN1, we next recorded spectra with the spin-label introduced at a site in between the pair of CAPRIN1 aromatic-rich regions (A658C, Fig. 2E,F) and next to the C-terminal arginine-rich segment (S678C, Fig. 2G,H). Notably, the *I*_*para*_ */ I*_*dia*_ profiles are strikingly similar for the three different spin-label positions for both the folded and unfolded states of E, E S-H pWT SOD1 (Fig. 2C-H), suggesting that the SOD1 interactions sites are broadly distributed across the CAPRIN1 sequence. The strong correspondence between *I*_*para*_ */ I*_*dia*_ values for the three CAPRIN1 cage sites strongly suggests that the placement of the cages does not interfere with intermolecular interactions in this system. To this end, the PRE experiments were repeated using a covalently attached nitroxide spin label, MTSL (43), with very similar *I*_*para*_ */ I*_*dia*_ profiles obtained relative to those from the DOTA cage.

As described above, the interactions of CAPRIN1 with folded E, E S-H pWT SOD1 were significantly weaker than with the unfolded ensemble. In order to probe CAPRIN1 contacts with the folded conformer more quantitatively we repeated the above experiments on a sample prepared with equimolar (1:1) ^15^N-labeled E, E S-H pWT SOD1 and spin-labeled ^14^N-CAPRIN1 (A658C, Fig. 2I). The resulting *I*_*para*_ */ I*_*dia*_ profile shows dips distributed across the sequence (Fig. 2I, *purple bars*) suggestive of potential specific interaction sites. However, these dips broadly overlap with sites that are predicted to be solvent-exposed and have high propensities for interaction, as inferred by the recognition factor amino acid scale (44) which assigns each amino acid a numerical value reflective of its hydrophilicity and average interaction energy with every other amino acid (Fig. 2I, *black circles vs. purple bars*). Indeed, mapping the intermolecular PRE profile onto the structure of a single protomer from the mature SOD1 structure (36) (PDB ID: 1HL5) indicates that the primary interaction sites are those that are most accessible (Fig. 2J).

### Conformational equilibrium of E, E S-H pWT SOD1 in CAPRIN1 condensed phase

Our NMR studies established that in a mixed solution comprised of E, E S-H pWT SOD1 and 3.5 mM CAPRIN1 the SOD1 folding – unfolding equilibrium shifts towards the unfolded state (from 80-95% folded, no CAPRIN1, to ~60% folded with 3.5 mM Caprin1, Fig. 1E). We wondered how the equilibrium would respond to higher concentrations of CAPRIN1, such as those found in the condensed phase of a demixed solution and how the structural dynamics of both folded and unfolded components of E, E S-H pWT SOD1 might differ between dilute and condensed phases. To this end we prepared a ^2^H, ^15^N, ^13^C-ILV labeled E, E S-H pWT SOD1 condensed phase sample following the protocol that is illustrated in *SI Appendix*, Fig. S3A. Starting from an initial solution of ∼250 µL of 1.85 mM E, E S-H pWT SOD1, in which a fractional population for the folded state of ~94% was quantified by NMR (*SI Appendix*, Fig. S3B, *purple*), 750 μL of unlabeled concentrated CAPRIN1 was added to a final concentration of 5.75 mM, reducing the folded population to ∼ 56% (*SI Appendix*, Fig. S3B, *green*). Phase separation was subsequently induced through the addition of NaCl until a final salt concentration of 150 mM was achieved. Note that the effect of CAPRIN1 addition results in a substantially greater reduction in the SOD1 folded population compared to the effect of 150 mM NaCl (*SI Appendix*, Table S1). After centrifugation of the phase-separated solution a bi-phasic mixture of dilute and condensed phases (*SI Appendix*, Fig. S3A) was produced and the mixture was transferred to a 3 mm NMR tube for studies by NMR. As the condensed phase occupies the entirety of the receiver coil only NMR signal originating from it is observed, while the dilute phase sitting above is NMR invisible (*SI Appendix*, Fig. S4A).

Prior to recording NMR spectra, we first measured the concentrations of the two major solvents in the condensed phase, CAPRIN1 and H_2_O. Given that both E, E S-H pWT SOD1 and CAPRIN1 are present in the sample, CAPRIN1 concentrations cannot be ascertained through absorbance measurements. Therefore, we prepared a reference CAPRIN1 condensed phase NMR sample, without SOD1, and measured the CAPRIN1 concentration via UV absorbance (28.3 mM). Then, by comparing signal integrals in both condensed phase samples described above, focussed over a region where exclusively CAPRIN1 resonances are found (1 – 2 ppm; note that SOD1 is deuterated with the ^1^H signals of ILV methyl groups resonating upfield of 1 ppm; *SI Appendix*, Fig. S4B, *green vs. purple*), the CAPRIN1 concentration in the CAPRIN1:E, E S-H pWT SOD1 condensed phase sample was determined to be 22.6 mM. Similarly, the relative H_2_O content in the condensed and dilute phases of the CAPRIN1:E, E S-H pWT SOD1 sample can be established from the ratio of H_2_O peak volumes in these two phases (V_H2O, Condensed_ / V_H2O, Dilute_) using pulse acquire measurements. We have performed experiments using flip angles for the water excitation pulse between 2.5o-45o, with a variety of receiver gains, and obtained consistent measures of 78±2 % for the relative water content in the condensed *vs*. the dilute phases (*SI Appendix*, Fig. S4C, *orange vs. blue*). These measurements establish that the CAPRIN1 concentration in the condensed phase is similar to that of water (CAPRIN1 ~245 mg/mL and H_2_O ~770 mg/mL) and, accordingly, CAPRIN1 plays a significant role in solvating SOD1.

To determine how the CAPRIN1 condensed phase solvent environment affects the folding equilibrium of E, E S-H pWT SOD1, we initially recorded [^1^H,^15^N] correlation spectra in both the dilute and condensed phases (Fig. 3A, *blue vs. orange*). Whereas peaks from both the folded and unfolded states of E, E S-H pWT SOD1 are observed in the dilute phase, only resonances derived from the unfolded ensemble are present in the condensed phase. Two possible scenarios can explain this result, including (i) that the CAPRIN1 condensed phase completely unfolds E, E S-H pWT SOD1 or (ii) that the slow tumbling of the folded conformer in the viscous CAPRIN1 condensed phase precludes observation of amide correlations. These two situations can be resolved by recording methyl-TROSY based spectra that can be an order of magnitude (or higher) more sensitive than amide-based experiments. To increase the sensitivity still further we have measured datasets using a delayed decoupling approach (45), referred to as ddHMQC (46), whereby one of the delays in the experiment is eliminated and the resulting time domain signal reconstructed to remove lineshape distortions that would otherwise result. Applications to the 1 MDa lumazine synthase complex showed very significant gains in signal-to-noise, including the appearance of crosspeaks that would otherwise not be observed in regular HMQC spectra. We have also modified the original ddHMQC experiment to include coherence transfer selection gradients for suppression of signals from the intense CAPRIN1 solvent, with careful design to minimize the length of the experiment (47). Unlike the situation with the [^1^H,^15^N] correlation experiment, where peaks derived from the folded state of E, E S-H pWT SOD1 were missing, all of the expected correlations from the folded E, E S-H pWT SOD1 conformer could be observed in the [^1^H,^13^C] ddHMQC spectrum of the condensed phase (Fig. 3B, *blue vs. orange*).

**Fig. 3.**
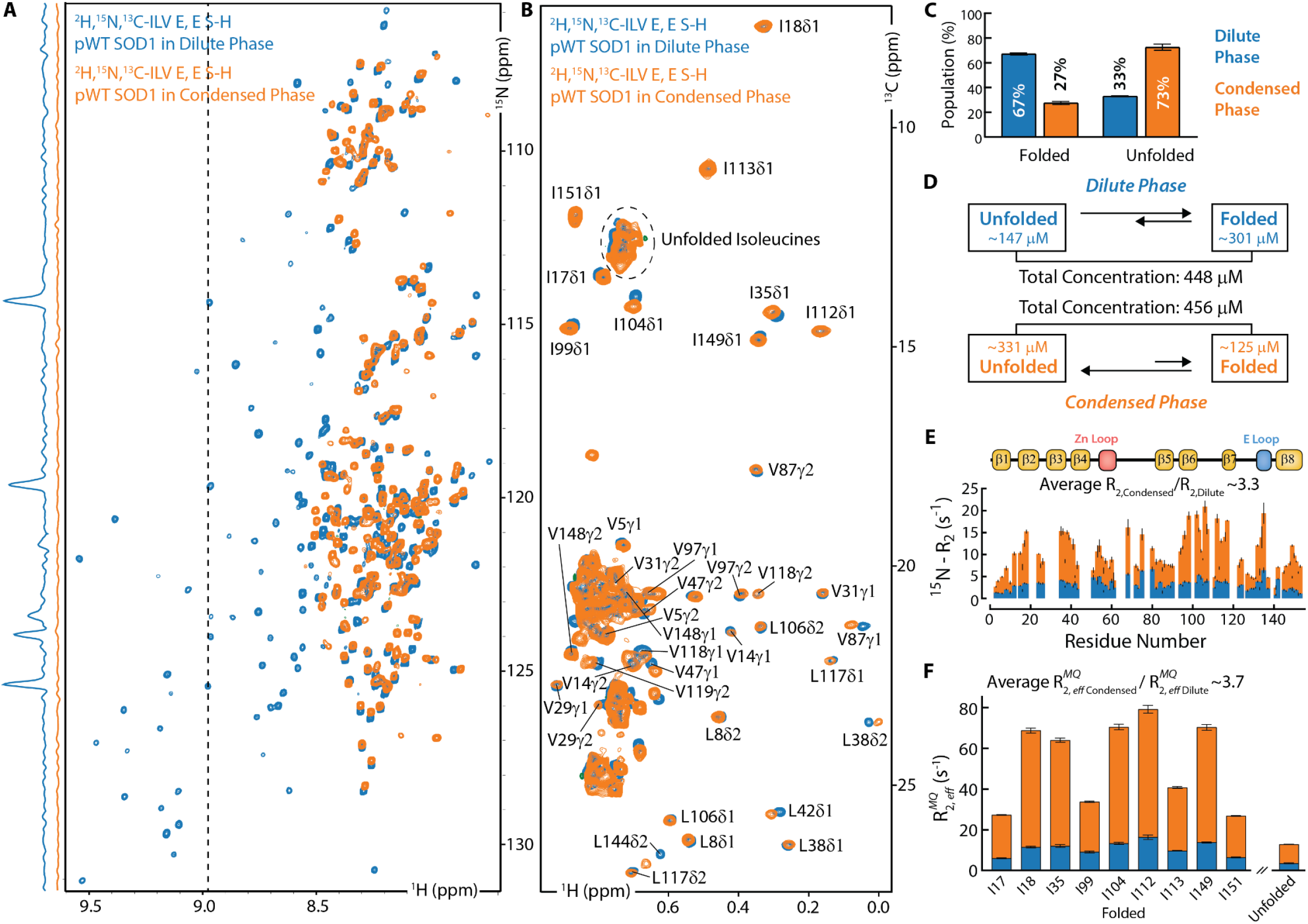
Effect of the CAPRIN1 condensed phase on the folding equilibrium and dynamics of E, E S-H pWT SOD1. (A)-(B) [^1^H, ^15^N] HSQC (A) and [^1^H, ^13^C] ddHMQC (B) spectra of ^2^H, ^15^N, ^13^C-ILV E, E S-H pWT SOD1 in dilute (*blue*) and condensed (*orange*) phases, 25 °C, 800 MHz. Vertical traces in A (left of 2D spectra) at the position of the dashed line illustrate that [^1^H, ^15^N] correlations from the folded state are observed in the dilute phase, but not in the condensed phase. In contrast, expected correlations for the folded conformer are observed in the ddHMQC spectra. (C) Quantification of folded and unfolded E, E S-H pWT SOD1 fractional populations based on integrals of resonances from folded and unfolded conformers recorded under fully relaxed conditions and taking into account the enhanced transverse relaxation rates of correlations in the condensed phase based on rates listed in panel F (*SI Appendix*, Materials and Methods). (D) Concentrations of folded and unfolded states of E, E S-H pWT SOD1 in the dilute (*blue*) and condensed (*orange*) phases from comparison to a reference spectrum with a known total concentration of protein. (E) Per-residue ^15^N *R*_2_ rates for the unfolded E, E S-H pWT SOD1 ensemble or resonances within disordered regions of the folded conformer in the dilute (*blue*) and condensed (*orange*) phases. Secondary structure elements of SOD1 are indicated above. (F) [^1^H-^13^C] multiple quantum relaxation rates of Ile δ1 methyl groups in folded E, E S-H pWT SOD1 in dilute (*blue*) and condensed (*orange*) phases. The average *R*_2_ increase reported pertains only to the folded isoleucine resonances. A similar ratio is reported for Ile δ1 methyl groups from the unfolded ensemble (right), however, only an average is obtained as the resolution in the current spectrum is not sufficient to resolve the correlations.

The high sensitivity of the [^1^H,^13^C] ddHMQC experiment enables a straightforward evaluation of how the E,E S-H SOD1 folding-unfolding equilibrium shifts in the condensed phase. To this end, peak volumes of Ile resonances from folded and unfolded states of E, E S-H SOD1 were quantified in a fully relaxed ddHMQC spectrum and compared with the corresponding volumes from a reference spectrum of SOD1 of known concentration. Corresponding measurements were made for the dilute phase of the demixed sample as well. In all cases peak volumes were corrected for transverse relaxation of signals during fixed delays in the experiment so that robust measures of concentrations of each state could be obtained (48). Notably, a decrease in the population of the folded conformer, from 67% in the dilute phase to 27% in the condensed phase, is observed (Fig. 3C), with concentrations of (unfolded, folded) SOD1 of ~147 µM, ~301 µM and of ~ 331 µM, ~125 µM in dilute and condensed phases, respectively (Fig. 3D). Note that while the CAPRIN1 condensed phase stabilizes the unfolded E, E S-H SOD1 ensemble, the total concentrations of SOD1 in the dilute and condensed phase are comparable (448 µM dilute phase *vs*. 456 µM condensed phase) differentiating the solvent impact on conformational equilibrium from partitioning.

The [^1^H,^13^C] ddHMQC spectrum of E, E S-H SOD1 in the condensed phase (Fig. 3B) clearly establishes that the highly concentrated CAPRIN1 solvent environment shifts the folding equilibrium of immature SOD1 towards the unfolded ensemble. However, the spectrum also indicates that the structure of the folded conformer in the condensate is unchanged from the conformation in the dilute phase (or buffer) as the positions of corresponding peaks are similar in both phases. We were interested in assessing whether the backbone and sidechain dynamics of E, E S-H SOD1 are also similar in the dilute and condensed environments. To this end we measured the ^15^N transverse spin relaxation rates for the backbone amides of the unfolded E, E S-H SOD1 ensemble in both phases (Fig. 3E). There is a large increase in rates in the condensed phase, as expected (compare blue *vs* orange, roughly 3-fold). The high amplitude segmental motions of the backbone chain, even in the condensed phase environment, are responsible for the excellent spectral quality of the unfolded state, and, correspondingly, the absence of such motions leads to the disappearance of resonances in amide correlation spectra derived from the folded E, E S-H SOD1 conformer. We also measured sidechain methyl [^1^H-^13^C] multiple quantum relaxation rates using a ddHMQC based Carr-Purcell-Meiboom-Gill experiment recorded with a small spacing between successive ^13^C refocusing pulses (CPMG frequency of 2 kHz) (47, 49), with relaxation rates increasing by approximately four-fold in the condensed phase (Fig. 3F). The increased motional degrees of freedom associated with the methyl-containing sidechains, coupled with methyl-TROSY based experiments, mitigate what would otherwise be very significant and, likely, pathological relaxation losses, highlighting the utility of methyl probes in studies of folded protein conformers in condensed phases.

Our relaxation data show that there is a significant attenuation of motion in both unfolded (backbone amides) and folded (sidechain methyls) E, E S-H SOD1. We wondered how translational diffusion would be affected by the condensed phase environment. *SI Appendix*, Fig. S4D shows signal attenuation profiles recorded using pulse field gradient-based diffusion experiments for E, E S-H SOD1 in both dilute (blue) and condensed (orange) phases using methyl group probes for the unfolded ensemble. Notably, the diffusion rate for unfolded E, E S-H SOD1 is ~ 8.5 times slower in the condensed phase relative to the dilute phase, 25 °C (*SI Appendix*, Fig. S4D, *orange vs. blue*), with E, E S-H SOD1 and CAPRIN1 diffusing with very similar rates in the condensate (1.04 × 10^−7^ cm^2^/s and 1.05 × 10^−7^ cm^2^/s). The signals for the folded state of SOD1 could not be observed during these relatively insensitive diffusion measurements.

### Unfolding of E, E S-H pWT SOD1 in the CAPRIN1 condensed phase is coupled to aggregation

The age-related decline in protein homeostasis and quality control mechanisms may contribute to the formation of aberrant condensates comprised of aggregated proteins (50). Given that a clinical hallmark of ALS is the presence of insoluble neuronal inclusions containing SOD1, among other proteins (31), we asked whether the condensed phase solvent environment could mediate this structural transition. To this end we monitored a CAPRIN1:E, E S-H pWT SOD1 condensed phase sample over a time period of approximately two months. Notably, we observed the formation of insoluble aggregates exclusively in the condensed phase fraction of our biphasic sample (Fig. 4A, *right vs. left*). ^1^H one-dimensional NMR measurements showed a relative decrease in signal intensities between spectra recorded at T = 0 and 62 days, indicating loss of soluble NMR-visible signals for both CAPRIN1 (Fig. 4B, *orange vs. black in gray shaded area*) and E, E S-H pWT SOD1 (Fig. 4B, *orange vs. black in purple shaded area*). By comparison, a CAPRIN1 condensed phase sample without SOD1 remained largely unchanged for a period of at least 20 months (Fig. 4C), while a sample of E, E S-H pWT SOD1 at a concentration comparable to the SOD1 concentration in the CAPRIN1:E, E S-H pWT SOD1 condensed phase was stable for at least 86 days (Fig. 4D). The relative stabilities of the different samples analyzed by NMR are summarized in Figure 4E. Negative-stain transmission electron microscopy (TEM) images of the CAPRIN1:E, E S-H pWT SOD1 condensed phase indicate that the aggregates formed adopt fibrillar morphologies (Fig. 4F). Note that aggregation is observed exclusively in the condensed phase via TEM despite comparable concentrations of total SOD1 in both phases of the phase-separated sample. Taken together our data are consistent with a prominent role for CAPRIN1 condensates in promoting the aggregation of the E, E S-H pWT SOD1 client via a coupled unfolding-aggregation mechanism.

**Fig. 4.**
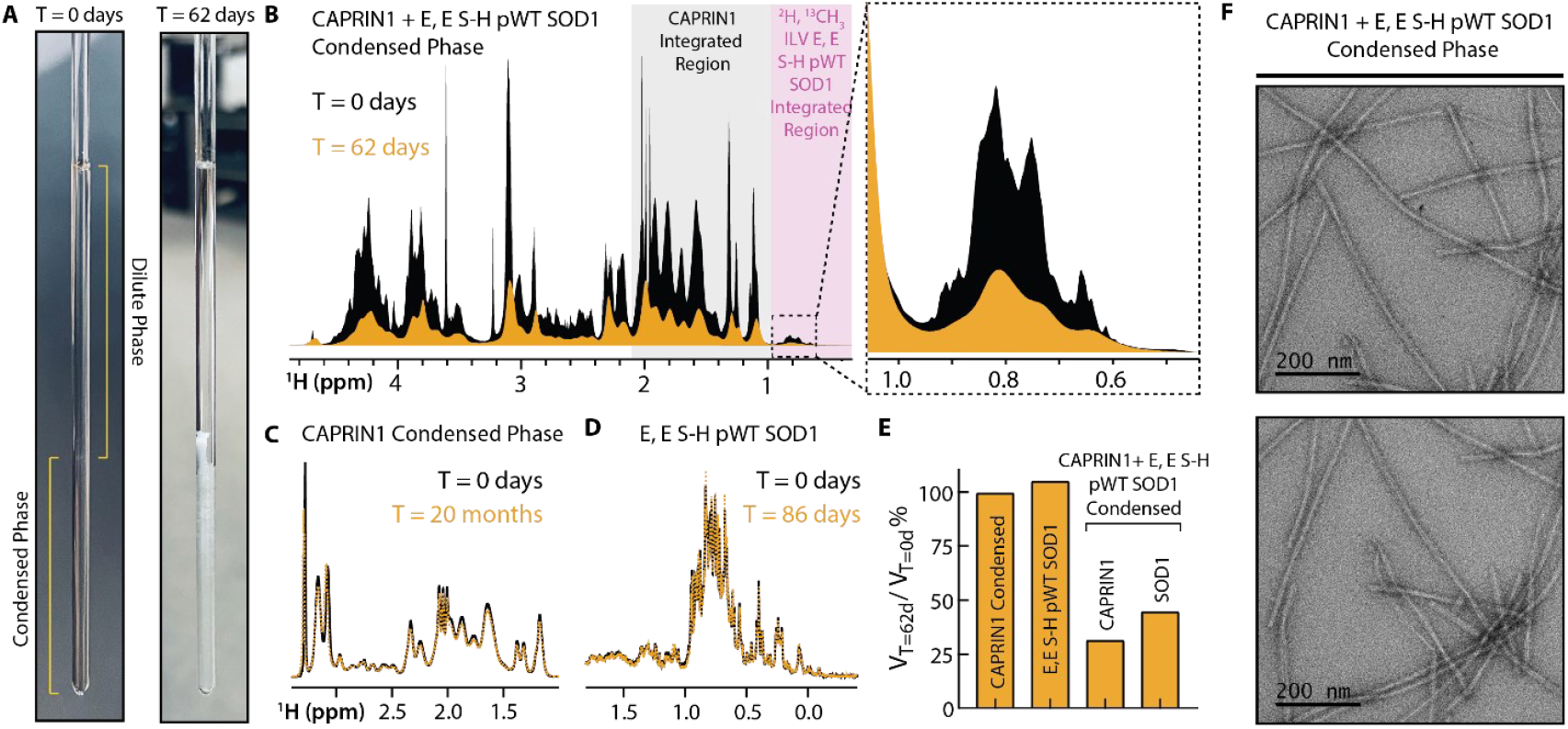
Coupled unfolding of SOD1 and aggregation of E, E S-H pWT SOD1:CAPRIN1 condensed phase. (A) A CAPRIN1: E, E S-H pWT SOD1 phase-separated sample at T=0 days (*left*) and at T=62 days (*right*), showing the formation of aggregates at the longer time point. (B) ^1^H NMR spectra of the sample at T=0 days (*black*) and at T=62 days (*orange*), indicating loss of soluble NMR-observable CAPRIN1 (*gray shaded region*; note that the variant of CAPRIN1 used does not have ILV residues) and loss of E, E S-H pWT SOD1 (*pink shaded region*). (C) ^1^H NMR spectrum of CAPRIN1 condensed phase (no SOD1) at T=0 (*black*) and at T=20 months (*orange dashed line*), indicating little change. (D) ^1^H NMR spectrum of 200 µM ^2^H, ^13^CH_3_-ILV SOD1 in NMR buffer at T=0 (*black*) and at T=86 days (*orange dashed line*). (E) Relative integrals of NMR-observable signals at T=62 days *vs*. T=0 days for CAPRIN1 in a CAPRIN1 condensed phase, E, E S-H pWT SOD1 in buffer, and CAPRIN1 and E, E S-H pWT SOD1 in a CAPRIN1:E, E S-H pWT condensed phase (*left to right*). The rate of change of the NMR observable signal is assumed to be linear between T=0 and the measured time point in the CAPRIN1 condensed phase and E, E S-H pWT SOD1 in buffer for computing the expected signal at T=62 days. (F) Negative-stain TEM images of the CAPRIN1:E, E S-H pWT SOD1 condensed phase diluted 100-fold with NMR buffer.

### The zinc-bound, dimeric form of SOD1 is less susceptible to unfolding than immature SOD1 when solvated by CAPRIN1

SOD1 maturation consists of a series of post-translational modifications, converting the nascent monomeric protein (T_m_ ~ 48 °C) into a thermodynamically stable (T_m_ ~ 92 °C) homodimer (28, 33, 34, 51). Binding of a single zinc ion to the high-affinity Zn-binding site (E, Zn S-H SOD1) is sufficient to drive partial dimerization of SOD1, restricting access to the immature state that would otherwise promote misfolding and aggregation (41). Subsequent copper-loading to the secondary metal-binding site and disulfide-formation (Cu, Zn S-S SOD1), catalyzed by the SOD1 copper chaperone, leads to an enzymatically active protein that is of high stability (51–53). We wondered whether the increased stability associated with the maturation process might protect against aggregation.

To this end we produced the zinc-bound, disulfide-reduced maturation intermediate (T_m_ ~ 58 °C) (33), Zn, Zn S-H pWT SOD1, where Zn is bound to the high-affinity metal binding site and predominately also to the second metal binding site which has a lower affinity for Zn ions. Under the conditions of our experiments (SOD1:Zn in the ratio 1:15), ~75% of SOD1 is calculated to exist in the Zn, Zn S-H SOD1_Dimer_ state (see *SI Appendix*), with the remaining (minor) states corresponding to the folded or unfolded conformers illustrated in Fig. 5A, *left*. We recorded ddHMQC spectra of pWT SOD1 with excess Zn both in the absence and presence of 3.5 mM CAPRIN1 (Fig. 5B, *green*), as was done for E, E S-H pWT SOD1 (Fig. 1D) in the absence of Zn (purple, dashed single line contours, Fig. 5B). Addition of CAPRIN1 generates new SOD1 resonances that overlap with signals from E, E S-H pWT SOD1 (purple), including those derived from both folded and unfolded states (Fig. 5B, *left vs*. Fig. 5B *right*). However, while the addition of CAPRIN1 shifts the distribution of SOD1 conformers to those less mature than Zn, Zn S-H pWT SOD1_Dimer_, such as Zn, Zn pWT S-H SOD1_Monomer_, for example, the fraction of folded SOD1 decreases only slightly, from 98% to 95% at the highest CAPRIN1 concentration added (3.5mM, Fig. 5C, *green*). This contrasts the situation for E, E S-H pWT SOD1 in the absence of Zn where addition of CAPRIN1 to 3.5 mM is accompanied by a large decrease in the population of the folded conformer, from 85% to 60% (Fig. 5C, *purple*) and concomitant increase in the amount of unfolded SOD1. Collectively, these findings establish that maturation of SOD1, such as formation of the zinc-bound, dimeric form, results in conformers that are more resilient to unfolding by CAPRIN1, underscoring the potential impact of ALS mutations that weaken metal binding or disrupt dimerization in terms of drivers of aggregation in condensates, leading possibly to disease.

**Fig. 5.**
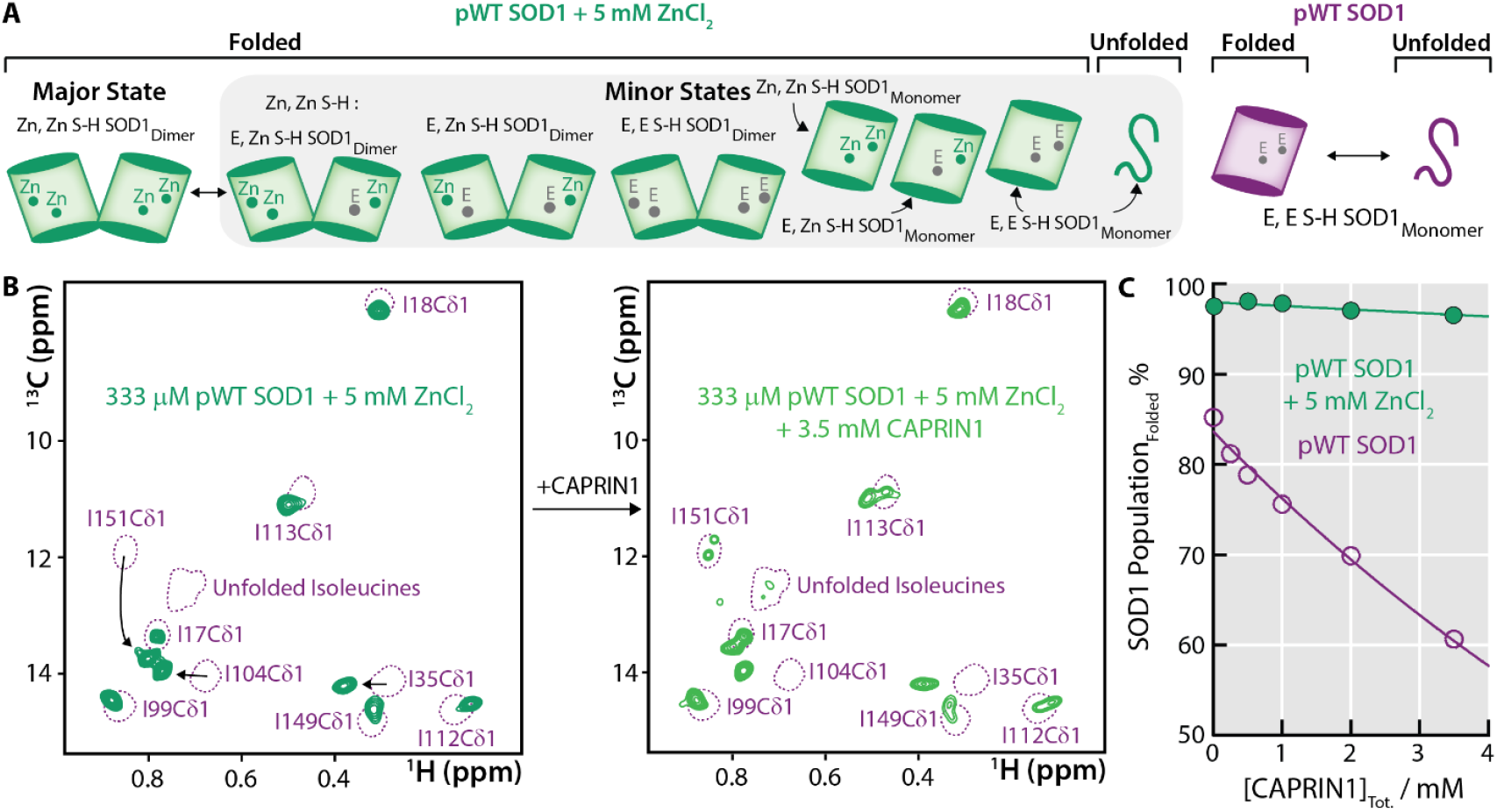
Metalated SOD1 is more resilient to unfolding when solvated by CAPRIN1. (A) Schematic indicating the conformational states of SOD1 produced in the absence of zinc or CAPRIN1 (*purple*) and in the presence of excess zinc (*green*). In the absence of zinc, SOD1 exists in equilibrium between folded and unfolded monomeric states (E, E S-H SOD1_monomer_). In the presence of excess zinc, SOD1 is primarily a folded dimer with full zinc occupancy of the primary, high-affinity, zinc binding site and high zinc occupancy at the secondary, low-affinity metal binding site (Zn, Zn S-H SOD1_Dimer_, ≈ 75%; see *SI Appendix*). Other minor states are shown in the diagram and are in equilibrium with the major zinc form. (B) Selected region of [^1^H,^13^C] ddHMQC spectra, 800 MHz, 25 °C, of 333 µM pWT SOD1 in the presence of 5 mM ZnCl_2_ (green), without (*left*) and with (*right*) 3.5 mM CAPRIN1. The corresponding ddHMQC spectra of pWT SOD1 (300 µM; *purple dashed*) is reproduced here from Fig. 1D for comparison. In the presence of CAPRIN1 (*right, green*), the populations of the minor states are elevated, giving rise to resonances that overlap with those from the metal-free pWT SOD1 state (*purple dashed*). (C) Population of pWT SOD1 folded conformations, including minor states, as a function of CAPRIN1 concentration. The metal-free pWT SOD1 *vs*. CAPRIN1 profile (purple) is reproduced from Fig. 1E for comparison.

## Discussion

Biophysical studies aim to provide a quantitative understanding of the kinetics and thermodynamics of interactions between biomolecules, a description of molecular structure at atomic resolution and how it influences these interactions, and insight into how structure, kinetics, and thermodynamics change over time in response to biological stimuli that elicit a particular function. Atomic resolution structural methods such as X-ray crystallography, electron cryo-microscopy, and NMR spectroscopy have all contributed to these efforts. However, these approaches have mainly focused on studies of folded and, in the case of NMR, also unfolded or disordered biomolecules in buffer solutions, and are currently largely unable to provide similarly rigorous descriptions of highly dynamic systems, particularly those associated with phase separation.

It is estimated that up to 80% of the components of the human proteome reside in biological condensates during some stage of their lifetimes (54) and the need to understand the structural dynamics of these molecules, and their nucleic acid counterparts, is thus critical. However, the highly dynamic nature of the interactions, including those between intrinsically disordered proteins and other protein molecules – both unfolded and folded – or nucleic acid elements, stymie X-ray and cryo-EM efforts. In principle, solution NMR spectroscopy offers a unique opportunity to provide an atomic description of the interactions that drive phase separation and how the condensed phase modulates biomolecular structural dynamics (10, 12, 15–17, 55–57). In practice, however, the high condensate viscosity broadens NMR spectra and places severe constraints on the types of experiments that can be performed (10, 15), especially for folded components for which signals from backbone amides in [^1^H,^15^N] HSQC-type spectra are, in general, not observed (Fig. 3A).

While there are difficulties associated with obtaining atomic resolution structural data on client molecules within condensates, a number of studies have generated indirect or regiospecific descriptions of how phase separation can regulate their function. For example, recent studies by the Rosen laboratory have shown that reaction rates can be accelerated in condensed phases, both by concentrating reaction components (mass action) and by molecular reorganization of these components that alters substrate *K*_*m*_ values (6). Relatedly, Gross and coworkers have shown that the activity of an mRNA decapping enzyme is enhanced through the addition of an activator in a condensed phase environment over what is observed in the dilute phase (24). In contrast, Arosio and coworkers have shown that condensates inhibit enzymatic activity of NADH-oxidase, despite elevated concentrations of reaction components in the condensed phase (58). Collectively, these findings suggest that the condensate solvent environment can remodel the free-energy landscapes of biomolecules, altering the conformations that are thermally accessible to them and, in turn, their function. Regiospecific insights on such protein structural changes in condensates have been reported, as inferred from hydrogen-deuterium exchange mass spectrometry measurements. The Sosnick laboratory has shown that different segments of the folded RNA-recognition motif domains of PolyA-binding protein 1 (Pab1) exhibit either increased or decreased deuterium uptake in Pab1 condensates relative to measurements in buffer, implying changes in folded domain stability in the condensed phase (59).

In principle, NMR studies of slowly tumbling structured molecules in highly viscous condensate environments are fraught with the same difficulties that hinder the investigation of molecular complexes with aggregate masses on the order of hundreds of kDa in buffer solutions. In the latter case the use of methyl-TROSY NMR (60), in concert with delayed-decoupling that minimizes signal intensity losses during the experiment (46, 47), has significantly alleviated the size limitations that would otherwise be pathologic. Here we have extended this approach to study the conformational changes of a “client” protein, E, E S-H pWT SOD1, inside a condensate comprised of the CAPRIN1 “scaffold”, at atomic resolution. Although peaks from the folded conformation of nascent SOD1 cannot be observed in [^1^H,^15^N]-based experiments, all of the expected methyl signals are visible in [^1^H,^13^C] ddHMQC spectra recorded using highly deuterated,^13^CH_3_ ILV-samples. Our results establish that the equilibrium between folded and unfolded E, E S-H pWT SOD1 conformations decreases from ~67% folded in the dilute phase to ~27% in the condensed phase of phase separated E, E S-H pWT SOD1:CAPRIN1 (Fig. 3C). Moreover, the degree of unfolding of SOD1 in mixed samples (*i.e*., absence of phase separation) titrates with the addition of CAPRIN1 (Fig 1E). PRE-based studies carried out in the mixed phase show that CAPRIN1 interactions with the SOD1 unfolded ensemble are more extensive than for the folded conformer, largely localizing to a region of unfolded SOD1 extending from residues H80-H120 (Fig. 2). CAPRIN1 displays multiple SOD1 interaction sites, as placement of spin labels at various positions along the CAPRIN1 chain resulted in attenuation of SOD1 signals (Fig. 2). The similarities between profiles of signal attenuation from the different spin label placements (Fig. 2C,E,G) suggest that CAPRIN1 interacts with the SOD1 client in a promiscuous manner, consistent with its solvent role. Our results also highlight that more mature forms of SOD1 are likely to be more stable in condensates, as observed for a doubly-zincated SOD1 molecule in a mixed phase environment that was highly enriched in CAPRIN1 (3.5 mM), with very little unfolding (Fig. 5), in contrast to nascent SOD1 (Fig. 1E). The correlation between unfolding and aggregation noted for E, E S-H pWT SOD1 (Fig. 4) suggests that mutations that impair metal binding and, hence stability, could play a role in aggregation in condensates, as observed in disease (32). It has been proposed that pathological aggregation in ribonucleoprotein granules arises from increases in local concentration of certain client proteins in these condensates (61, 62). Our results suggest a more nuanced perspective wherein, aggregation can occur without concentrating the client, via shifts in the conformational equilibria towards aggregation-prone states.

This study also highlights the important role that CAPRIN1, and by extension the proteinaceous environment in condensed phases in general, plays in acting as a solvent of client molecules. Indeed, our measurements indicate that the concentrations of CAPRIN1 (245 mg/mL) and water (770 mg/mL) are comparable in the condensed phase (*SI Appendix*, Fig. S4), substantiating the role of CAPRIN1, and other scaffolds, as solvents. Further, we demonstrate that the physical characteristics of the proteinaceous solvent, such as their net charge, govern the free-energy landscape of client molecules (Fig. 1F). Given that cells dynamically alter condensate composition through PTMs, these findings underscore a mechanism by which the solvent nature of condensates can be modulated, and, accordingly the conformational states of resident proteins, to achieve specific biological functions.

Our work points to the unique role that solution NMR spectroscopy can play in quantifying changes to free energy landscapes driven by highly concentrated protein environments, and in providing atomic resolution insights into the mechanism by which these changes occur. The information available from such studies is complementary to the exciting results forthcoming from other biophysical studies (25, 63) and together will provide a more detailed understanding of the way in which phase separation regulates cellular function.

## Acknowledgements

R.A., A.K.R., and S.K.H. acknowledge post-doctoral support from the Canadian Institutes of Health Research (CIHR). J.D.F.-K. acknowledges support from CIHR (FDN-148375, PJT-190060) and from the Canada Research Chairs Program. L.E.K. acknowledges support from the CIHR (FND-503573) and the Natural Sciences and Engineering Council of Canada (2015-04347).

## Notes

### Competing Interest Statement

The authors have declared no competing interest.

