## Supplementary Text, Figures, Tables and Materials and Methods for "Atomic resolution map of the solvent interactions driving SOD1 unfolding in CAPRIN1 condensates"

**SI Text for "*The zinc-bound, dimeric form of SOD1 is less susceptible to unfolding than immature SOD1 when solvated by CAPRIN1*"**

Consider the following sequential binding model for zinc binding to SOD1 (1),

$$A + B \overset{K_1}{\leftrightharpoons} AB$$

$$AB + B \overset{K_2}{\leftrightharpoons} AB_2 \qquad [S1]$$

where $A$ refers to E, E S-H pWT SOD1 and $B$ to $Zn^{2+}$, and the association (dissociation) constants for $Zn^{2+}$ binding to the Zn and Cu sites are $K_1$ ($K_{d1}$) and $K_2$ ($K_{d2}$), respectively.

It has been shown previously that (1),

$$a[AB]^3 + b[AB]^2 + c[AB] + d = 0$$

where

$$a = 4K_{d1} - K_{d2},$$

$$b = 4K_{d1}K_{d2} - {K_{d2}}^2 + 2K_{d2}A_T,$$

$$c = K_{d1}{K_{d2}}^2 + {K_{d2}}^2A_T + {K_{d2}}^2B_T + K_{d2}{B_T}^2 - 2K_{d2}A_TB_T,$$

$$d = -{K_{d2}}^2A_TB_T.$$

For a total concentration of E, E S-H pWT SOD1, $[A_T]$, of 333 μM and a total $Zn^{2+}$ concentration, $[B_T]$, of 5,000 μM, and for $K_{d1} \leq$ 0.1 μM and $K_{d2}$ = 80 μM (1), respectively, the fraction of E, Zn S-H pWT SOD1 is 1.8% and Zn, Zn S-H pWT SOD1 is 98.2%, with a negligible population of E, E S-H pWT SOD1. Further consider the dimerization equilibrium $2M \leftrightharpoons D_2$ where $M$ is the pWT SOD1 monomer and $D_2$ the dimer, with a dissociation constant of $K_{d,dimer}$. It follows that:

$$[M] = \frac{-K_{d,dimer} + \sqrt{(K_{d,dimer})^2 + 8K_{d,dimer}P_T}}{4} \qquad [S2]$$

where $P_T = [M] + 2[D_2]$ is the total protein concentration. At a total protein concentration of 333 μM and a $K_{d,dimer}$ of 51 μM for SOD1 with at least one metal binding site bound to zinc (1), the fraction of SOD1 protomers that are in the dimeric state is 76%. Thus, under our experimental conditions, ~75% of SOD1 exists in the Zn, Zn S-H pWT $SOD1_{Dimer}$ state upon addition of 15-fold excess of Zn.

### Materials and Methods

#### Expression and purification of CAPRIN1

The C-terminal low-complexity disordered region of human CAPRIN1 comprising residues Ser607-Gln707 and harboring the mutations N623T, N630T, V610A, L621A, referred to as CAPRIN1 in what follows, was subcloned into a pET-His-SUMO vector and transformed into *Escherichia coli* BL21 (DE3) RIPL cells. The cells were grown to an $OD_{600}$ ~ 0.6 – 0.8, induced with 0.5 mM IPTG and expressed overnight at 25 °C in LB. Cells were harvested via centrifugation, resuspended in lysis buffer (6 M guanidinium hydrochloride, 50 mM Tris pH 8, 500 mM NaCl, 10 mM Imidazole), and sonicated for 15 minutes (2 s on, 2 s off). Lysed cells were spun down at 40,000 g for 45 minutes, and the supernatant was loaded onto a Ni-NTA column (GE Healthcare) equilibrated with lysis buffer. Following extensive washing with lysis buffer, CAPRIN1 was eluted with 50 mM Tris pH 8, 150 mM NaCl, 400 mM imidazole. Cleavage of the His-SUMO tag was achieved through overnight dialysis at 4 °C into 50 mM Tris pH 8, 150 mM NaCl, 10 mM imidazole, 2 mM β-mercaptoethanol in the presence of HisSUMO protease. The cleaved protein was subjected to a Ni-NTA column to remove the His-SUMO tag and HisSUMO protease, concentrated, and loaded onto a Superdex75 (26/600) column equilibrated with 3 M guanidinium hydrochloride, 50 mM Tris pH 8, 500 μM EDTA. The purified protein was stored at -20 °C until use.

**Expression, purification, and preparation of various SOD1 states**

The human SOD1 2-154 construct containing the C6S and C111A mutations, referred to as pseudo-WT (pWT) SOD1, and an N-terminal His tag was transformed into *Escherichia coli* BL21 (DE3) RIPL cells. The cells were grown to an $OD_{600}$ ~ 0.6-0.8, induced with 0.5 mM IPTG, and expressed overnight at 20 °C in M9 $D_2O$ ($H_2O$) media supplemented with 3 g/L $d_7$-glucose (glucose) for deuterated (non-deuterated) protein production. Incorporation of $^{19}F$ on the tryptophan 32 sidechain was achieved by growing cells in M9 $H_2O$ media supplemented with 120 mg/L of 5-fluoro D/L-tryptophan, 60 mg/L of L-phenylalanine, 60 mg/L of L-tyrosine and 1 g/L of glyphosate one hour prior to induction of protein expression (2). Uniform $^{15}N$ isotopic labeling was achieved by the addition of 1 g/L $^{15}N$ ammonium chloride to the M9 growth medium. Methyl labeling was achieved by the addition of 100 mg/L of 2-keto-3-methyl-$d_3$-3-$d_1$-4-$^{13}C$-butyrate (for non-sterospecific labeling of Leu, Val-$^{13}CH_3$/$^{12}CD_3$) and 60 mg/L of 2-keto-3-$d_2$-4-$^{13}C$-butyrate (for labeling of Ileδ1) one hour prior to the induction of protein expression (3).

Cells were harvested by centrifugation, resuspended in lysis buffer (6 M guanidinium hydrochloride, 20 mM HEPES pH 8, 10 mM Imidazole), and sonicated for 15 minutes (2 s on, 2 s off). Lysed cells were spun down at 40,000 g for 45 minutes, and the supernatant was loaded onto a Ni-NTA column (GE Healthcare). After extensive washing with lysis buffer, SOD1 was refolded on the column by washing with non-denaturing buffer (20 mM HEPES pH 8, 10 mM imidazole, 20% glycerol) and eluted with 20 mM HEPES pH 8, 400 mM imidazole, 20% glycerol. Cleavage of the His tag was achieved through overnight dialysis at room temperature into 50 mM HEPES pH 8, 2 mM DTT in the presence of HisTEV protease. The cleaved protein was isolated through another round of Ni-NTA purification, concentrated, and loaded onto a Superdex75 (26,600) column equilibrated with 20 mM HEPES pH 8. The pure protein-containing fractions were pooled and stored at 4 °C until use (typically within 3-4 days).

The E, E S-H pWT $SOD1_{Monomer}$ (immature state) was prepared by first removing bound metals through overnight dialysis at 4 °C into 50 mM sodium acetate and 50 mM EDTA pH 3.82. The protein was then subjected to another round of dialysis into Milli-Q water for 6 hours, and subsequently concentrated to ~ 5 mL. An equivalent volume of denaturing buffer (8 M guanidinium hydrochloride, 40 mM HEPES pH 7.4, 2 mM EDTA), pre-treated with Chelex 100 resin (Sigma 95621) and filtered, was added to the SOD1 in Milli-Q water and the mixture was kept at room temperature for 30 minutes. Freshly Chelexed and filtered TCEP was added to the mixture at a final concentration of 10 mM and allowed to react for 3 hours at 37 °C to reduce disulfide bonds. The mixture was injected into a HiPrep 26,10

Desalting column pre-equilibrated with Chelex 100, filtered NMR Buffer (20 mM HEPES pH 6.3, 1 mM TCEP, 1 mM $NaN_3$, 10% $D_2O$), and the protein-containing fractions were pooled, and concentrated as needed. SOD1 is refolded during this buffer exchange step, resulting in a folded population that varies from prep-to-prep in the range of ~80-95%. Note that comparisons in the absence *vs.* presence of CAPRIN1 in mixed solutions and in dilute *vs.* condensed phases are made using the same SOD1 stocks, such that the change in populations are a reflection of the CAPRIN1 solvent environment.

The Zn, Zn S-H pWT $SOD1_{Dimer}$ and associated minor states (Fig. 5A) were prepared by adding $ZnCl_2$ at a final concentration of 5 mM to 500 µM E, E S-H $SOD1_{Monomer}$ in NMR buffer, prepared as described above. The mixture was allowed to incubate for 30 minutes at room temperature. Subsequently, the SOD1 was diluted to a final concentration of 333 µM with NMR buffer containing 5 mM $ZnCl_2$ and/or CAPRIN1 in the same buffer. Under these conditions, the population of Zn, Zn S-H $SOD1_{Dimer}$ is ~75% (see above), with additional folded minor states formed with complete, partial, and no incorporation of zinc, both in the dimeric and monomeric forms, as well as an unfolded non-metallated state (Fig. 5A).

**Preparation of CAPRIN1: E,E S-H pWT $SOD1_{Monomer}$ condensed phase**

The CAPRIN1:immature SOD1 condensed phase was prepared as shown schematically in *SI Appendix*, Fig. S3A. CAPRIN1 was buffer-exchanged into Chelexed, filtered NMR buffer (20 mM HEPES pH 6.3, 1 mM TCEP, 1 mM $NaN_3$, 10% $D_2O$), using a a HiPrep 26,10 Desalting column. Concentrated stocks of CAPRIN1 (> 7 mM) and $^{2}H$, $^{15}N$, $^{13}C$-ILV E, E S-H pWT SOD1 (> 1.5 mM) were mixed on ice and allowed to equilibrate for 30 minutes, generating the mixed phase solutions shown in Fig. S3 (*green*). Phase separation was induced by the addition of NaCl (150 mM final concentration). The phase-separated mixture was centrifuged at 21,000 g for 3 minutes using a microcentrifuge, producing a two-phase system (dilute phase on top, condensed phase on bottom). The mixture was transferred to a 3-mm NMR tube that had been pre-treated overnight with concentrated nitric acid and washed extensively with MilliQ water. The condensed phase volume was sufficient to occupy the entirety of the receiver coil with the dilute phase sitting above, as shown in *SI Appendix*, Fig. S4A. After completion of condensed phase NMR measurements, some of the dilute phase sitting above was decanted into a separate 3-mm NMR tube to generate the dilute phase sample used for measurements reported in Fig. 3.

**Incorporation of Dia- and Paramagnetic metal-bound cages onto CAPRIN1 and measurement of phase separation propensity**

Maleimido-mono-amide-DOTA cage was purchased from Macrocyclics (B-272) in powdered form and resuspended in 4 M guanidinium hydrochloride, 50 mM Tris pH 7 buffer to a final concentration of 100 mM. Cysteines were introduced into CAPRIN1 in the N623T, N630T, V610A, and L621A mutant background with the following additional mutations: S615C, A658C, and S678C (one cysteine mutation at a time; CAPRIN1 607-707 has no endogenous cysteines). A 5-fold molar excess of the DOTA cage was added to each of the CAPRIN1 cysteine mutants in 4 M guanidinium hydrochloride, 50 mM Tris pH 7, 1 mM TCEP, and allowed to react overnight at 37 °C. Reaction completion was confirmed by mass spectrometry. The DOTA-cage-bound-CAPRIN1 cysteine mutants were each divided into two aliquots of equal volume, and either gadolinium (III) chloride or lutetium (III) chloride was added in 50-fold molar excess. Following overnight incubation at 37 °C, the reaction mixtures were buffer exchanged using a HiPrep 26,10 Desalting column pre-equilibrated with Chelex 100, filtered NMR buffer (as above), to remove excess DOTA cage and metals.

Measurement of water $T_1$s confirmed comparable DOTA-cage coordination of gadolinium for the three different cysteine mutants. Phase separation propensity of the cage-bound CAPRIN1 was determined using $A_{600}$ nm measurements at a constant [CAPRIN1] of 400 μM and in the presence of increasing [NaCl] up to a final [NaCl] of 2 M (*SI Appendix*, Fig. S2), using a BioPhotometer D30 (Eppendorf), error bars were obtained as the standard deviation of triplicate measurements.

**Negative stain transmission electron microscopy of CAPRIN1:immature SOD1 condensed phase** (*Fig. 4F*)

Continuous carbon grids were prepared using 400 mesh copper/rhodium grids (Electron Microscopy Sciences) by applying a layer of collodion support on the rhodium side. A ~4 nm carbon film was deposited on the collodion support using a Leica EM ACE200 carbon thread evaporation coater under pulse mode. Grids were glow-discharged in air for 15 s immediately prior to sample application. The CAPRIN1:immature SOD1 condensed phase sample (22.6 mM; T = 62 days) was diluted ~100-fold with NMR buffer and vortexed extensively prior to application on the carbon grids. 4 μL of the sample was applied to each grid and allowed to adsorb for 2 minutes. The grid was rinsed by touching the sample side of the grid to a series of drops of milli-Q water (for a total of three times, wicking away excess water between each rinse), stained with 2% uranyl acetate for 10 s, and allowed to air-dry after wicking away excess stain. Samples were imaged with an FEI Tecnai F20 electron microscope equipped with a 200 keV field emission gun and applying a defocus range of 2 to 3 μm. Images were acquired with a Gatan Orius SC1000A camera with a 10 s exposure, using a nominal magnification of 25,000x, corresponding to a pixel size of 2.65 Å.

**NMR measurements**

All $^{1}$H, [$^{1}$H,$^{15}$N], and [$^{1}$H,$^{13}$C] NMR spectra were acquired either on an 18.8 Tesla (800-MHz $^{1}$H frequency) Bruker Avance III HD spectrometer or a 23.5 Tesla (1 GHz $^{1}$H frequency) Bruker Avance Neo spectrometer equipped with cryogenically cooled x, y, z pulsed-field gradient triple-resonance probes. All $^{19}$F-detected NMR measurements were performed at 11.7 Tesla (500 MHz $^{1}$H frequency) on a Bruker III HD spectrometer equipped with a liquid nitrogen-cooled z pulsed-field gradient triple-resonance probe. Spectra were processed using NMRPipe (4) and analyzed with either NMRPipe, peakipy (https://github.com/j-brady/peakipy) or NMRFAM-SPARKY (5).

*[$^{1}$H, $^{13}$C] methyl-TROSY ddHMQC NMR measurements of SOD1 folded and unfolded populations in mixed solutions (Fig. 1E)*

Methyl-TROSY spectra were recorded at 25 °C 800 MHz with a [$^{1}$H, $^{13}$C] ddHMQC pulse scheme with gradients for coherence transfer selection (6) for quantification of SOD1 folded and unfolded populations. Spectra were acquired under fully relaxed conditions with an interscan delay of 5 s, 4 scans/FID, $^{13}$C spectral widths, and acquisition times of 21 ppm and 10 ms, respectively, and a total acquisition time of 31 minutes/spectrum. $^{1}$H pulses and $^{13}$C pulses were centered on the water line and 19 ppm, respectively. Water suppression was obtained using gradients for coherence transfer selection; a water-selective EBURP1 pulse (7) (~ 7 ms) is applied at the outset to ensure that the water signal is along the +Z axis at the start of $^{1}$H acquisition.

Integrals of the nine well-dispersed folded isoleucine resonances (there are 9 Ile in SOD1) and the overlapping unfolded cluster were measured and the folded population fraction ($P_F$) was calculated as:

$$P_F = \frac{\sum V_i^F}{\sum V_i^F + \sum V_i^U} \quad [S3]$$

where $V_i^F$ is the volume of the "folded" (F) peak *i*, and $V_i^U$ is the volume of the "unfolded" (U) peak *i*. The volume of the unfolded isoleucine cluster was determined through a sum-over-box approach.

*$^{19}$F NMR for measurement of SOD1 folded and unfolded populations in mixed solutions (Fig. 1F and Fig. S1)*

$^{19}$F NMR spectra of the tryptophan 32 sidechain resonance of E, E S-H pWT SOD1 (only a single Trp in the protein) in the absence and presence of a number of crowders were acquired with an anti-ringing triple pulse excitation scheme to relieve baseline distortions, as described previously (8), at 25 °C on a 500 MHz spectrometer. Spectra were recorded with an interscan delay of 5 s, 10,240 scans, and $^{19}$F spectral widths and acquisition times of 50 ppm and 87 ms, respectively. $^{19}$F pulses were centered at -125 ppm.

[$^1$H,*$^{15}$N*] *TROSY HSQC NMR for measurement of intermolecular CAPRIN1:E, E S-H pWT SOD1 interactions via the paramagnetic relaxation enhancement effect (Fig. 2)*

[$^1$H,$^{15}$N] TROSY HSQC spectra (9) were recorded on samples of $^{15}$N E, E S-H pWT SOD1 in the presence of CAPRIN1 conjugated with either a Gd-bound DOTA cage (paramagnetic) or Lu-bound DOTA cage (diamagnetic). Spectra were acquired at 25 °C on a 1 GHz spectrometer with an interscan delay of 5.5s required for >99% recovery of Z-magnetization, determined from integrals of the SOD1 NH spectral region as a function of interscan delay using a diamagnetic sample. [$^1$H,$^{15}$N] TROSY HSQC spectra were acquired with 4 scans/FID, $^1$H and $^{15}$N spectral widths and acquisition times of 16.1 ppm and 26.5 ppm, and 64 ms and 50 ms, respectively, for a total acquisition time of 1 hour and 42 minutes/spectrum. $^1$H and $^{15}$N pulses were centered on the water line and 119 ppm, respectively. The paramagnetic relaxation enhancement effect was quantified by taking the ratio of corresponding peak intensities in spectra recorded on the paramagnetic and diamagnetic samples ($I_{para}$ / $I_{dia}$). Error bars were derived from the standard deviation of replicate measures.

*Measurement of CAPRIN1 concentration in the CAPRIN1:E, E S-H pWT SOD1 condensed phase via $^1$H NMR (SI Appendix, Fig. S4B)*

The concentration of CAPRIN1 in the CAPRIN1:E, E S-H pWT SOD1 condensed phase was determined by comparing peak integrals with a reference CAPRIN1 condensed phase sample (no SOD1) of known concentration, as illustrated in *SI Appendix,* Figure S4B. Spectral regions from 1-2.1 ppm were compared, containing only peaks from CAPRIN1 (note that SOD1 is deuterated, with the exception of ILV methyl groups and thus does not contribute to signals in this region). Excitation-sculpting $^1$H NMR spectra (10) were recorded at 25 °C on an 800 MHz spectrometer with an interscan delay of 5 s, 128 scans, and spectral width and acquisition times of 16 ppm and 64 ms, respectively. The concentration of CAPRIN1 in the condensed phase of the reference sample was determined by $A_{280}$ measurements (extinction coefficient: 10, 430 $M^{-1}cm^{-1}$) of a 2 μL aliquot, diluted 100-fold with 8M guanidinium chloride to generate a well-dispersed solution amenable to absorbance measurements.

*Measurement of $H_2O$ content in the dilute and condensed phases of phase-separated samples (Fig. S4C)*

The relative amount of $H_2O$ in the condensed and dilute phases was determined based on the ratio of $H_2O$ peak integrals ($V_{H2O, Condensed}$ / $V_{H2O, Dilute}$) in $^1$H NMR spectra. Single-scan spectra were recorded at 25 $^o$C on an 800 MHz spectrometer using a simple pulse acquire scheme with a low receiver gain to prevent receiver overflow, and an acquisition time of 64 ms. $^1$H pulses were centered on the water line, and a series of spectra were recorded with flip angles of {45, 22.5, 10, 5, 2.5} and subsequently quantified. The fraction of water in the condensed phase as compared to the dilute phase ranged between 75.5 and 79.6% (average of 77.6 ± 1.5%, where 1.5% is one standard deviation in the range of quantified values). Spectra were also recorded with a number of different receiver gains, with similar water ratios obtained.

*Single quantum and triple quantum pulse-field gradient (PFG) NMR for measurement of SOD1 diffusion constants in the dilute and condensed phases (SI Appendix, Fig. S4D)*

Diffusion measurements of $^2$H, $^{15}$N, $^{13}$C-ILV E, E S-H pWT SOD1 in the dilute phase were carried out using a single quantum, stimulated-echo based pulse scheme described previously (11), with the exception that $^{15}$N and $^{13}$C pulses were interchanged. Diffusion constants were obtained by integrating SOD1 methyl $^1$H signals in 1D experiments, recorded as a function of encoding /decoding gradient strengths, each applied as a bipolar gradient pair. The length of each of the encoding and decoding gradients was 0.9 ms for a total time δ of 1.8 ms, with a diffusion delay (Δ) of 150 ms. Condensed phase diffusion constants of E, E S-H SOD1 were obtained using a triple quantum-based pulse scheme (12), with Δ = 150 ms and lengths of encoding/decoding gradients (δ) totaling 3.23 ms. Diffusion measurements were recorded at 25 $^o$C on a Bruker 800 MHz spectrometer (maximum gradient strength of 44.6 G/cm).

*[$^1$H, $^{15}$N] correlation spectra and $^{15}$N-$R_{1\rho}$, $^{15}$N-$R_1$ measurements for probing the dynamics of unfolded SOD1 and disordered regions of folded SOD1 in CAPRIN1 dilute and condensed phases (Fig. 3A, E)*

[$^1$H,$^{15}$N] HSQC spectra of $^2$H, $^{15}$N, $^{13}$C-ILV E, E S-H pWT SOD1 in both the dilute and condensed phases of CAPRIN1 were recorded in the gradient, enhanced sensitivity mode (13) at 25 $^o$C, 800 MHz. Spectra were recorded with an interscan delay of 1.5 s, 8 scans/FID, $^1$H and $^{15}$N spectral widths and acquisition times of 15.0 ppm and 26.5 ppm, and 64 ms and 45 ms, respectively, for a total acquisition time of 43 minutes/spectrum. $^1$H and $^{15}$N pulses were centered on the water line and 119 ppm, respectively.

$^{15}$N $R_1$ and $R_{1\rho}$ rates of $^2$H, $^{15}$N, $^{13}$C-ILV E, E S-H pWT SOD1 were recorded in both dilute and condensed phases of CAPRIN1 using gradient enhanced sensitivity-based HSQC experiments, 25 $^o$C 800 MHz (14). Condensed (dilute) phase $R_{1\rho}$ measurements were collected with a series of six time points extending from 5 ms - 85 ms (5 ms – 100 ms). Six time points ranging from 10 ms - 700 ms were obtained for measurements of $R_1$ in both dilute and condensed phases.

$$R_2 = (R_{1\rho} - R_1\cos^2\theta) / \sin^2\theta \qquad [S4]$$

where θ = arctan(v1/ΔΩ), v1 is the spin-lock field strength (2 kHz), and ΔΩ is the offset (Hz) of the spin in question from the $^{15}N$ carrier (15).

*[$^{1}H$, 13C] methyl-TROSY ddHMQC NMR for measurement of SOD1 folded and unfolded populations in CAPRIN1 dilute and condensed phases of phase-separated samples (Fig. 3B-D)*

Methyl-TROSY spectra were recorded on dilute and condensed phase regions of phase separated samples, 25 °C, 800 MHz, using a [$^{1}H$,$^{13}C$] ddHMQC pulse scheme with gradients for coherence transfer selection (6). Spectra were acquired with an interscan delay of 1.5 s, 32 scans/FID, $^{13}C$ spectral widths, and acquisition times of 21 ppm and 25 ms, respectively, and a total acquisition time of 3 hours and 4 minutes/spectrum. $^{1}H$ and $^{13}C$ pulses were centered on the water line and 19 ppm, respectively. Stereospecific assignments of the leucine and valine methyl resonances were obtained from an analysis of the relative phase of the methyl resonances in a [$^{1}H$,$^{13}C$] CT-HSQC spectrum recorded on a fractionally (10%) $^{13}C$-labeled SOD1 sample, as described previously (16).

For quantification of folded and unfolded E, E S-H pWT SOD1 populations, the ddHMQC-based methyl-TROSY experiments described above were recorded under fully relaxed conditions with an interscan delay of 5 s, 16 scans/FID, a $^{13}C$ acquisition time of 18 ms and a spectral acquisition time of 2 hours and 12 minutes. $P_F$ and $P_U$ were determined from the sum of the integrals of the folded ($V_i^F$) and unfolded ($V_i^U$) isoleucine resonances, accounting for differences in transverse relaxation for the folded *vs*. unfolded species, as follows:

$$P_F = \frac{\sum \frac{V_i^F}{e^{-R_{2,eff,i}^{MQ} * t_{seq}}}}{\sum \frac{V_i^F}{e^{-R_{2,eff,i}^{MQ} * t_{seq}}} + \sum \frac{V_i^U}{e^{-R_{2,eff,i}^{MQ} * t_{seq}}}} \qquad \text{[S5]}$$

where $V_i^F$ is the volume of "folded" (F) peak *i*, and $V_i^U$ is the volume of "unfolded" (U) peak *i*, $t_{seq}$ is the length of all fixed delays during the [$^{1}H$,$^{13}C$] ddHMQC experiment (4.6 ms), and $R_{2,eff,i}^{MQ}$ is the intrinsic methyl multiple quantum transverse relaxation rate for peak *i*. $V_i^U$ was estimated using a box-sum approach.

*ddHMQC-based CPMG NMR experiments for measurement of methyl multiple quantum transverse relaxation rates of SOD1 in dilute and condensed phases (Fig. 3F)*

Methyl multiple quantum transverse relaxation rates ($R_{2,eff}^{MQ}$) were acquired at a static magnetic field of 800 MHz, 25 °C, using a ddHMQC-based Carr-Purcell-Meiboom-Gill (CPMG) experiment (6, 17) with a constant-time CPMG element ($T_{relax}$) of 40 ms and 10 ms in dilute and condensed phases, respectively. Spectra were acquired with an interscan delay of 2 s and, $^{13}C$ acquisition time and spectral width of 18 ms and 21 ppm, respectively. Dilute (condensed) phase spectra were recorded with 24 (32) scans/FID. $^{1}H$ pulses and $^{13}C$ pulses were centered on the water line and 19 ppm, respectively. CPMG profiles, $R_{2,\,eff} = -1/T_{relax}\ln(I/I_0)$ *vs*. $\nu_{CPMG}$, were generated from peak intensities with ($I$) and without ($I_0$) the CPMG relaxation element of duration $T_{relax}$. For the dilute phase, 18 $\nu_{CPMG}$ points were recorded, ranging from 25 Hz to 2,000 Hz. Given that only flat profiles were observed for all of the resonances, the average (standard deviation) of the $R_{2,eff}^{MQ}$ values across the 18 $\nu_{CPMG}$ points was used as a measure of the intrinsic multiple quantum relaxation rate (error). For the condensed phase, the CPMG experiment was reduced to a two-point analysis recorded with ($I$) and without ($I_o$) a $\nu_{CPMG}$ pulsing element (2 kHz). Error values are computed from S/N of the peaks.

**Table S1. SOD1 folded and unfolded populations in different solutions.**

| SOD1 Solutions | [NaCl] / mM | [CAPRIN1] / mM | Folded Population % | Unfolded Population % |
|---|---|---|---|---|
| NMR Buffer | 0 | 0 | 93.5 | 6.5 |
| NMR Salt Buffer | 150 | 0 | 81.2 | 18.8 |
| Mixed Solution | 0 | 5.75 | 56.3 | 43.7 |
| Dilute Phase | 150* | 0.42 | 67.1 | 32.9 |
| Condensed Phase | 150* | 22.6 | 27.5 | 72.5 |

*The bulk concentration of NaCl added to induce phase separation is reported.

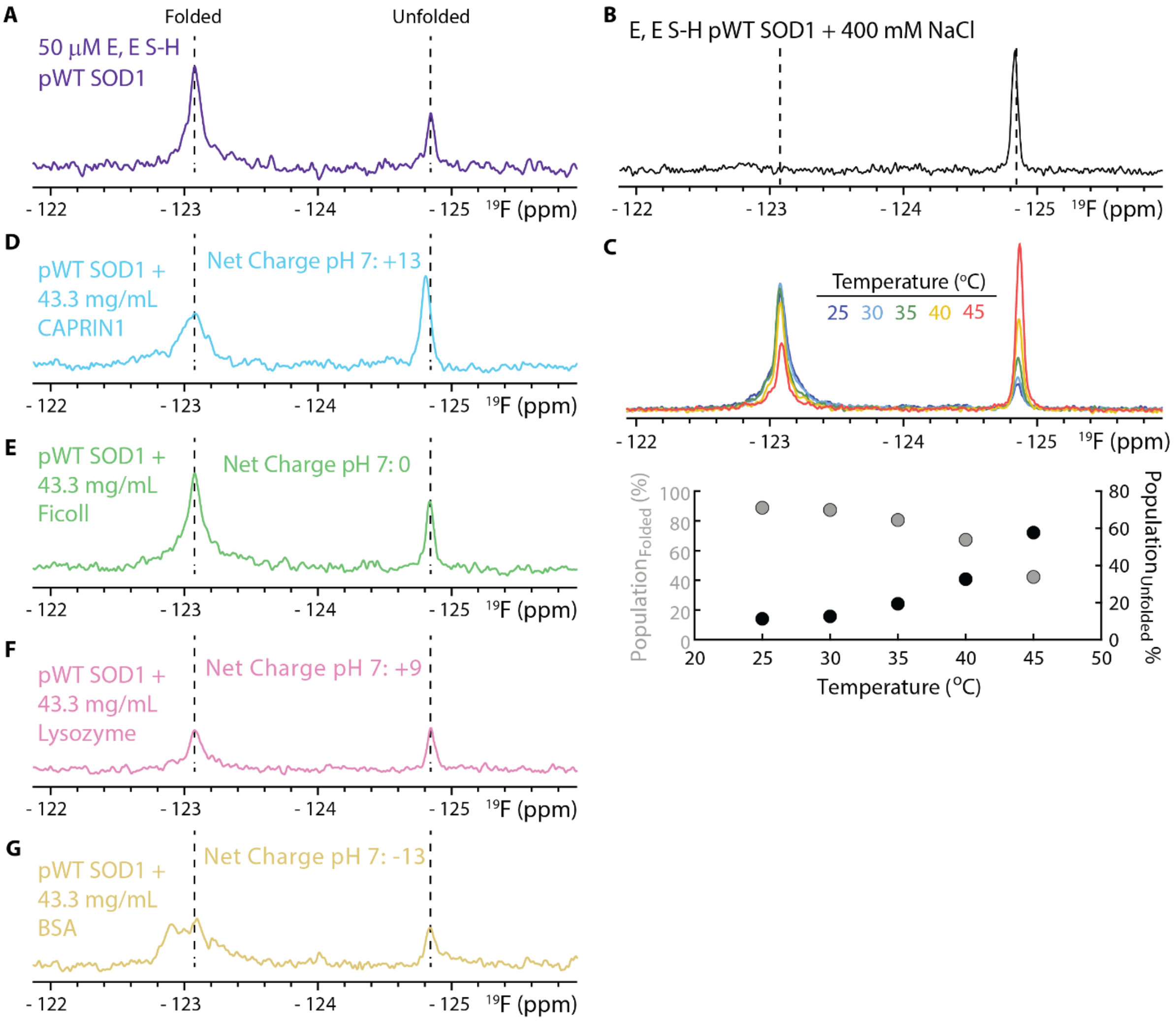


**Fig. S1 Effect of solvent environment on the folding equilibrium of E, E S-H pWT SOD1.** (A) $^{19}$F NMR spectrum of E, E S-H pWT SOD1 labeled at position W32 with 5-fluorotryptophan, 25 °C, 500 MHz. Two separate peaks corresponding to folded and unfolded SOD1 are annotated, assigned on the basis of salt (B) and temperature (C) measurements (small changes in $^{19}$F peak positions with temperature were "corrected" based on a temperature titration of 5-fluorotryptophan); high salt and high temperature stabilize the unfolded state. (D) – (G) $^{19}$F NMR spectra of 50 µM E, E S-H pWT SOD1 in the presence of 43.3 mg/mL of (D) CAPRIN1, (E) ficoll, (F) lysozyme, and (G) BSA. Addition of the crowders, in particular lysozyme, resulted in some precipitation, leading to a decrease in SOD1 protein concentration and consequently peak volumes in NMR spectra.

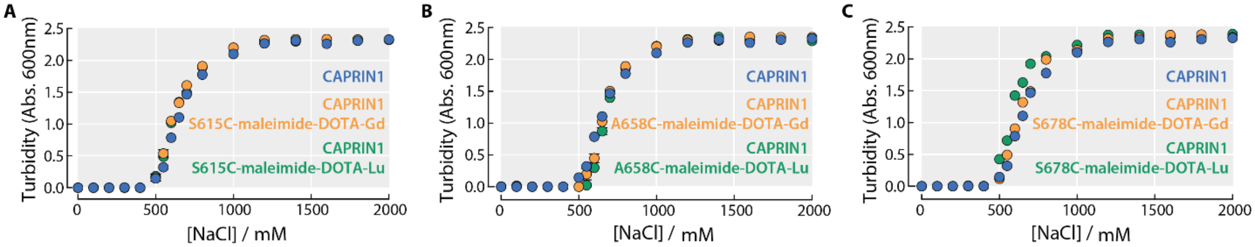


**Fig. S2 Metal-bound cages do not compromise CAPRIN1 phase-separation propensities.** (A) Salt-dependent phase separation of 400 μM CAPRIN1, as probed through changes in solution turbidity, for unmodified CAPRIN1 (*blue*), and CAPRIN1 modified with either a maleimide-DOTA-Gd cage (*orange*) or a maleimide-DOTA-Lu cage (*green*) attached to an introduced cysteine at position 615. Error bars are based on the standard deviation of replicates (smaller than most data points). (B) – (C) As (A), except for a cage introduced on a cysteine at position 658 (B) or 678 (C). All experiments were recorded at 25 °C.

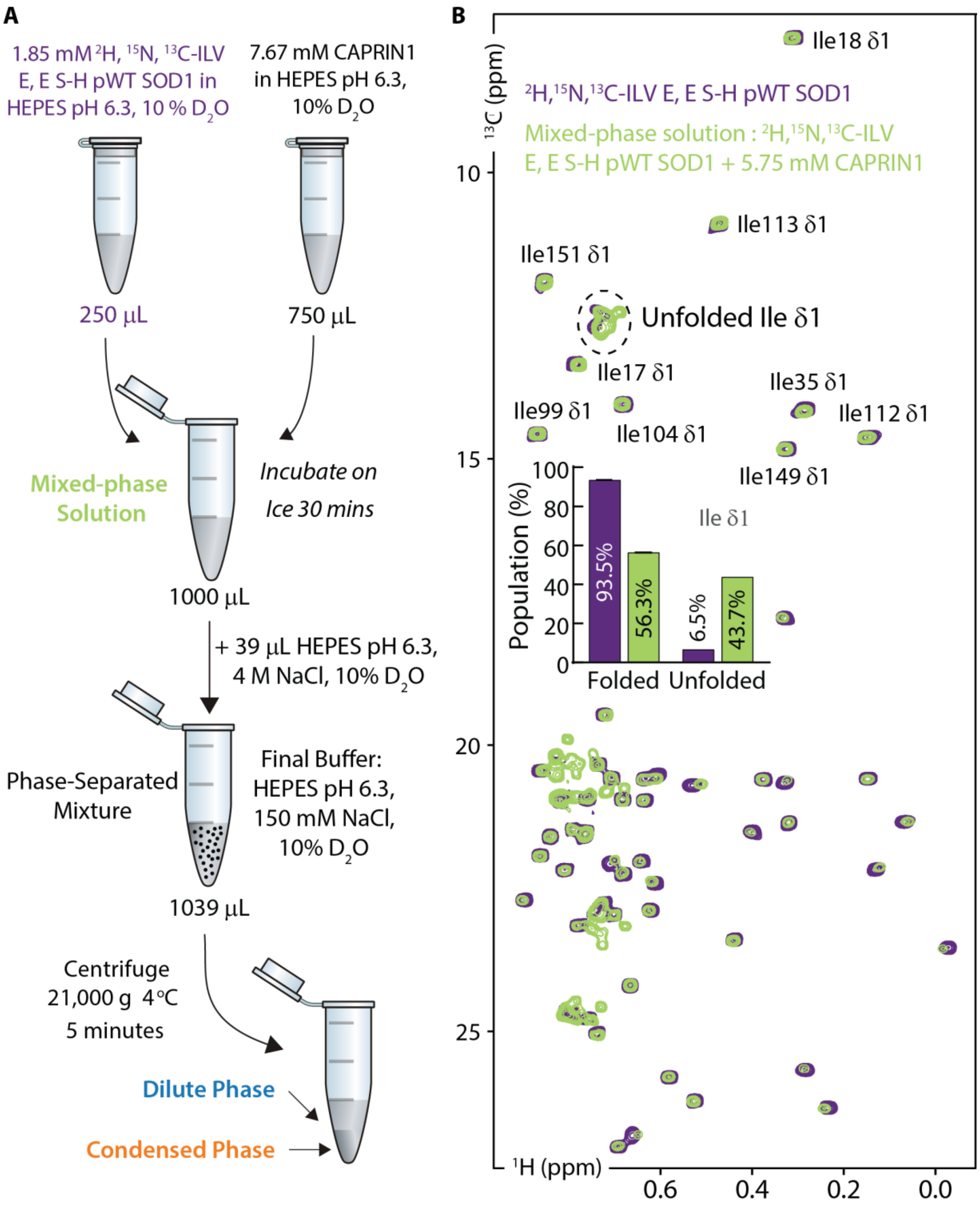


**Fig. S3 Preparation of dilute and condensed phases of CAPRIN1:E, E S-H pWT SOD1 and measurement of SOD1 folding–unfolding equilibrium at different stages of the sample preparation protocol.** (A) Schematic depicting the preparation of CAPRIN1:E, E S-H pWT SOD1 dilute and condensed phase samples. Sample labels are color-coded in correspondence with the respective spectra recorded. (B) [$^1$H,$^{13}$C] ddHMQC spectra of 200 μM, $^2$H, $^{15}$N, $^{13}$C-ILV E, E S-H pWT SOD1, no NaCl (*purple*) and a mixed-phase solution containing $^2$H,$^{15}$N,$^{13}$C-ILV E, E S-H pWT SOD1 and 5.75 mM CAPRIN1, no NaCl (*green*), 25 °C, 800 MHz. The inset shows the populations of folded and unfolded states based on integrals of Ile δ1 methyl resonances.

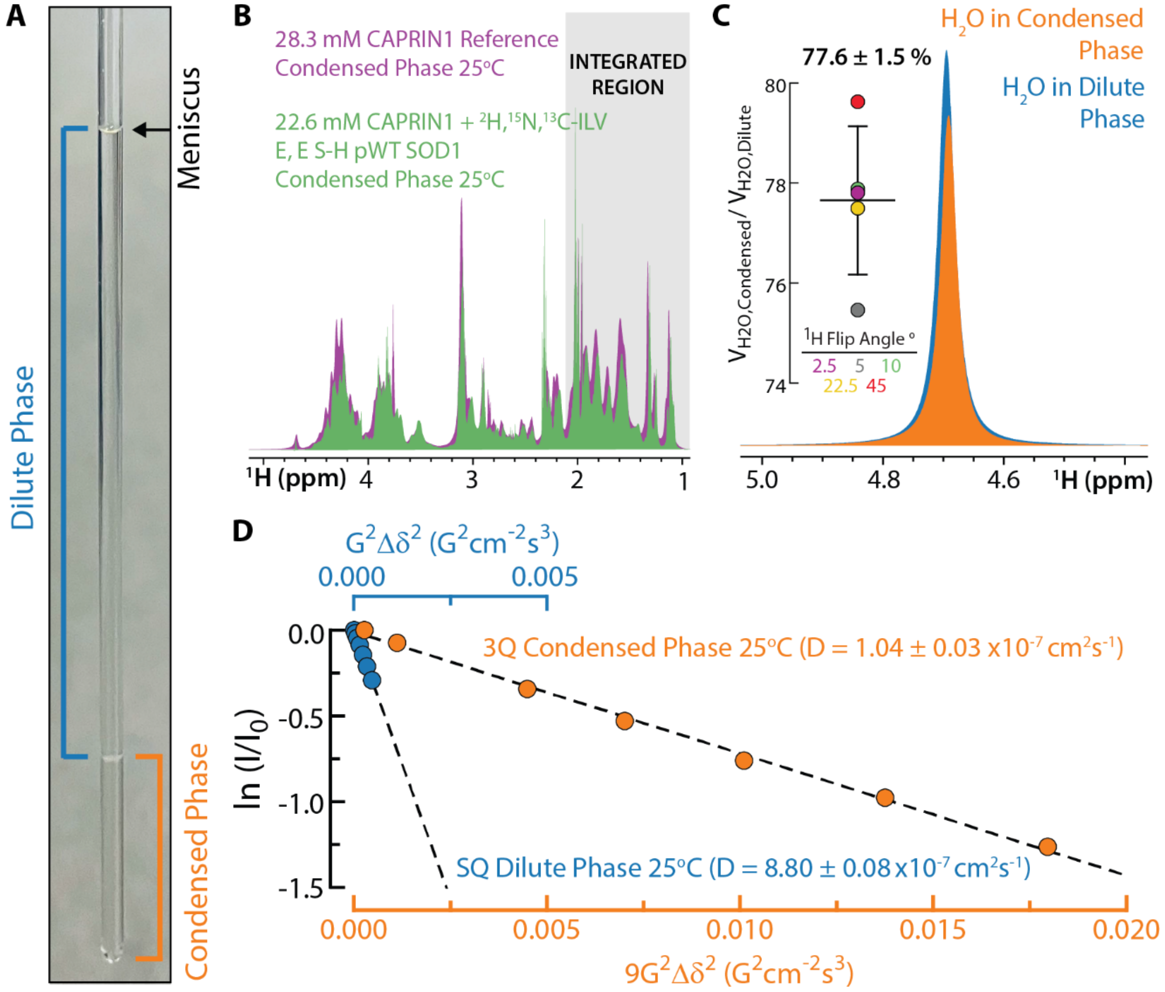


**Fig. S4 Material properties of the CAPRIN1:E, E S-H pWT SOD1 condensed phase.** (A) The CAPRIN1:E, E S-H pWT SOD1 phase-separated sample in a 3mm NMR tube. The condensed phase (*orange*) occupies the entire length of the receiver coil, with the dilute phase (*blue*) above. A separate sample of the dilute phase alone (*SI Appendix*, Materials and Methods) is used for measurements reporting on the dilute phase. (B) Overlay of $^{1}H$ NMR spectra of a reference CAPRIN1 condensed phase sample (*purple*) of known concentration (28.3 mM based on $A_{280}$ nm measurements, *SI Appendix*, Materials and Methods) and the CAPRIN1:E, E S-H pWT SOD1 condensed phase (*green*). A comparison of integrals of the aliphatic region (*grey shaded area*), not including resonances upfield of 1 ppm where SOD1 ILV peaks are found, was used to estimate the concentration of CAPRIN1 in the CAPRIN1:E, E S-H pWT SOD1 condensed phase. (C) Superposition of water signals in $^{1}H$ NMR spectra of dilute (*blue*) and condensed (*orange*) phases comprising CAPRIN1 and E, E S-H pWT SOD1, 25 °C. Spectra were acquired with a flip angle of 45°, and quantification of the relative peak volumes at several flip angles are shown in the panel inset. (D) Measurement of unfolded E, E S-H pWT SOD1 diffusion constants (25 °C) in dilute (*blue*) and condensed (*orange*) phases using single (SQ; dilute phase, (11)) and triple (3Q; condensed phase, (12)) quantum based pulse field gradient experiments. The experimental data (points) were fit (dashed lines) to obtain diffusion coefficients, as described in *SI Appendix*, Materials and Methods.
